# Epistasis limits but does not prevent the transfer of mutation-drug resistance mapping across 600 million years of fungal evolution

**DOI:** 10.64898/2026.07.08.737038

**Authors:** Marie-Ève Picard, Romain Durand, Alexandre K Dubé, Soham Dibyachintan, Alicia Pageau, Philippe C Després, Emilie Alexander, Jordan Grenier, Rong Shi, Christian R Landry

## Abstract

Whether different pathogens acquire resistance to antimicrobials through the same mutations is a major question in evolution and microbiology. Most antifungal drugs are used to treat infections caused by multiple fungal species, many of which have diverged for millions of years. If the evolution of resistance was to converge onto the same set of mutations across species, knowing the mechanism of resistance in one would allow us to predict and track it in others. The extent of this convergence remains unknown due to the lack of systematic data on resistance mutations. Here, we quantify the conservation of resistance mutations in the cytosine deaminase (CD), a protein responsible for resistance to flucytosine, one of the oldest antifungal drugs. By comparing the crystal structures of this enzyme through 600 My of evolution, we show that the CD structure is highly conserved. We compared the full CD mutational resistance spectrum of resistance from an ascomycete and a basidiomycete. We found that resistance mutations in one ortholog can be used to predict resistance in the other at a high level of accuracy. However, because of epistasis, around 10% of mutations have distinct effects in the two orthologs, which imposes an upper limit to the transferability of the knowledge of resistance mutations from one species to another. Using biochemical assays and by structural characterization of several mutants, we identify distinct mechanisms of epistasis, one important being that the local physiochemical environment of some position has evolved in a way that makes the same substitutions destabilizing or entirely inactivating in an ortholog-specific manner. Our results show that resistance mutations can be conserved in fungi across hundreds of millions of years of evolution but that epistasis eventually limits the accuracy of these predictions.

## Introduction

The same mutations often affect protein function differently when introduced into orthologous genes^1–3^. As proteins diverge, they accumulate amino acid differences that may not affect protein function but alter intramolecular interactions, thereby changing which amino acid is evolutionarily tolerated at a given site^4^. This type of interaction, also called epistasis, can occur locally or at a distance through allostery. Unexpected differences in the outcome of mutations can come from specific epistasis, whereby changes in the protein sequence modulate the effect of only particular amino acid changes. They can also result from global epistasis whereby mutations affect general biochemical properties such as expression, stability or activity, in such a way that the phenotypic outcome of many mutations will differ in one homolog compared to the other. From a biomedical point of view, both types of epistasis have numerous consequences, including on the predictability of the impact of mutations in proteins and organismal phenotypes^5^.

Being able to predict the impact of mutations on resistance is particularly relevant in the context of antimicrobial drug resistance, where information on mutational effects in one species could be used to infer resistance phenotypes in another. Indeed, antimicrobial drugs are often used to control the growth of distinct species sharing orthologs of the same protein target. For instance, the same antifungal drugs are used to treat infections caused by dozens of fungal species that have diverged for millions of years^6^. Due to epistasis, mutations that cause resistance in one species may have no or even opposite phenotypic impact when introduced at the same position in another species. This suggests that the wealth of data that have accumulated on model fungal species^7,8^, pathogens and non pathogens, would be of limited use to interpret genomic variation in other species, including emerging pathogens. High transferability would drastically reduce the efforts required to estimate the impact on resistance from any mutation in a pathogenic organism, i.e., an organism often lacking crucial genetic editing tools to quickly test mutational effects in the lab. Although there is currently no complete data that would allow us to assess the importance of this phenomenon quantitatively for antifungal drugs, some results suggest on antibiotic resistance that epistasis significantly alters the transferability of the effect of mutations from one ortholog to another^9,10^. For instance, a recent study on bacterial B1 metallo-β-lactamases found substantial variation in mutational effects across two homologs diverging at ∼70% of protein residues, with almost a third of mutations at conserved positions exhibiting homolog-dependent effects^9^.

Fungi acquire antifungal drug resistance through a common set of genes^11^. One of them is the cytosine deaminase (CD, encoded by *FCY1*) involved in the pyrimidine salvage pathway. CD is a small homodimeric protein that canonically converts cytosine into uracil^12^ and also converts the antifungal 5-fluorocytosine (5-FC) to 5-fluorouracil (5-FU), which makes it of broader medical interest as 5-FU is used as an anticancer drug^13^. As for many other genes involved in antifungal resistance^7^, mutations conferring resistance in CD are loss of function (LOF)^7,14^. A systematic assessment of the impact of all amino acid substitutions in CD from *Saccharomyces cerevisiae* (ScCD) revealed that resistance-conferring mutations are enriched at the dimer interaction interface and at buried positions, implying that protein destabilization is a major mechanism of resistance^14,15^. Nearly one third of all amino acid substitutions assayed resulted in partial or complete LOF, indicating that a large fraction of spontaneous mutations in *FCY1* can confer resistance. This raises the question whether resistant mutations in ScCD also confer resistance in CD homologs from fungal pathogens (Fig. 1a). Measuring the overlap between the mutational landscapes of resistance of ScCD and a CD homolog should provide invaluable insight into the extent of epistasis and whether or not it prevents predicting resistance using data from homologs.

**Figure 1:**
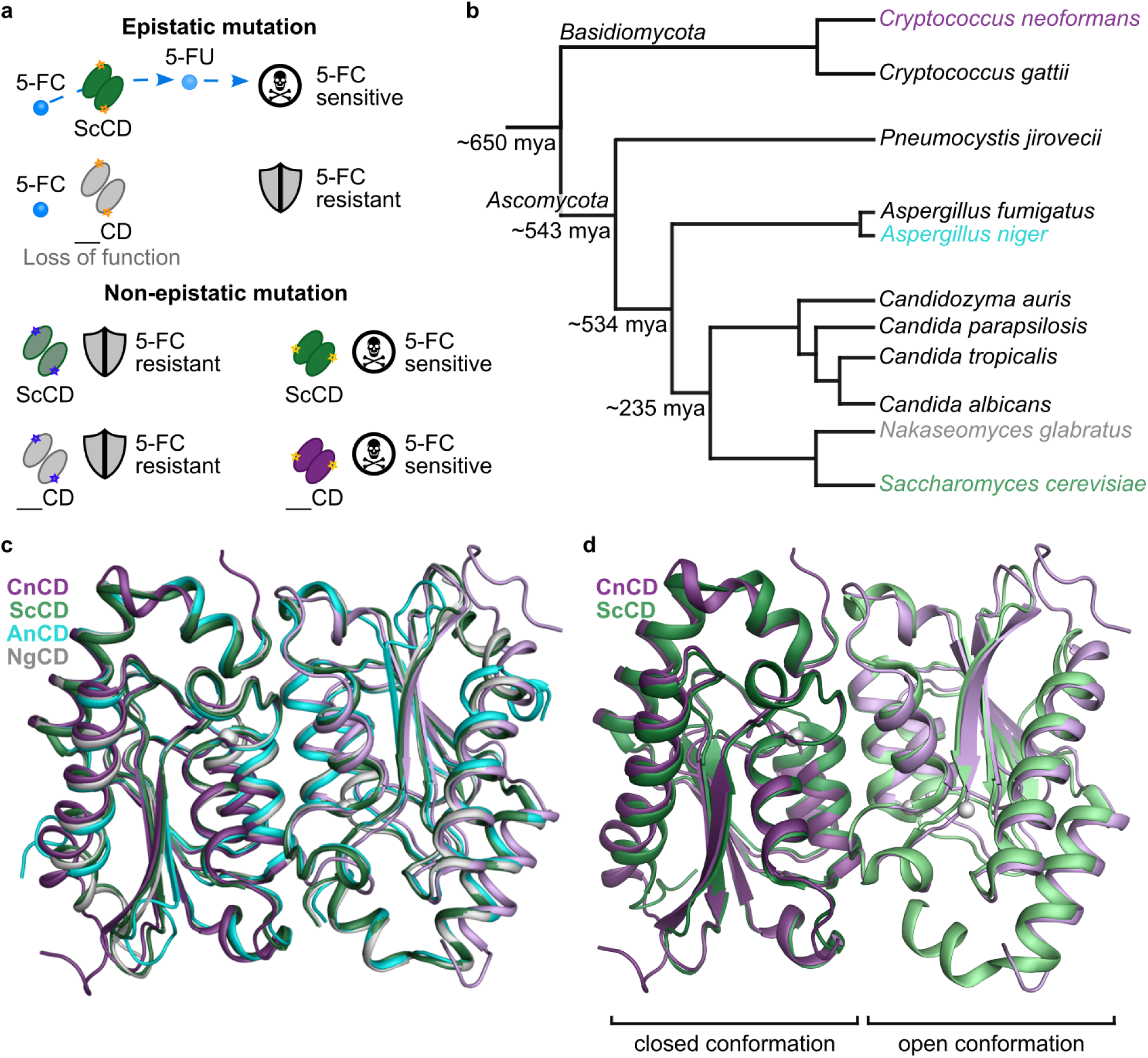
Conservation and divergence in the fungal cytosine deaminase (CD). (**a**) Simplified mode of action of 5-fluorocytosine (5-FC) and the implications of epistasis. 5-FC is a prodrug, which after import is deaminated to 5-fluorouracil (5-FU) by the cytosine deaminase (CD). 5-FU is toxic, which means a loss of function in CD is sufficient for survival (5-FC resistance). Whenever the same mutation has a different effect when introduced in CD orthologs, the mutation is said to be modulated by epistasis. On the other hand, if the same mutation has a comparable effect in both backgrounds (either loss of function leading to 5-FC resistance, or the mutation is 5-FC sensitive in both backgrounds), it is labeled non-epistatic. The “skull and bones” icon was created by Dmitry Vasiliev from Noun Project. (**b**) Simplified phylogenetic tree of pathogenic fungal species across *Ascomycota* and *Basidiomycota*^19^. Time of divergence obtained from TimeTree^20^. (**c**) Structural alignment of CD orthologs from *C. neoformans* (CnCD in purple, PDB 36FA), *Aspergillus niger* (AnCD in cyan, PDB 36FB), *Nakaseomyces glabratus* (NgCD in gray, PDB 36FC) and *S. cerevisiae* (ScCD in green, PDB 8VLJ). Structures of CnCD, AnCD, and NgCD were experimentally determined in this study. For all but CnCD, both subunits are in the closed conformation. For CnCD, the subunit on the left is the closed conformation, while the one on the right (colored a lighter shade of purple) is the open conformation. (**d**) Structural alignment of open-closed ScCD (green, PDB 8VLK: subunits A and B) and open-closed CnCD (purple, PDB 36FA). Panels (**c**) and (**d**) were generated in PyMOL v 3.1.0^21^, with zinc ions represented as spheres.

The most common clinical use of 5-FC is to treat cryptococcal meningitis (in combination with liposomal amphotericin B), an acute infection caused by several *Cryptococcus* species^16^. Resistance to 5-FC in this species occurs through LOF mutations in the pyrimidine salvage pathway, including in *Cryptococcus neoformans* CD (CnCD)^17,18^. Comparing a handful of mutations introduced in ScCD and CnCD using *S. cerevisiae* for heterologous expression showed that the phenotypic consequences are broadly conserved, i.e., most mutations caused comparable phenotypes in both orthologs^14^. This is a promising observation given the evolutionary distance between ScCD and CnCD. Indeed, this would suggest that such conserved mutational effects would behave the same way in any fungal pathogen whose CD homolog has diverged more recently (Fig. 1b). However, 9 out of 27 mutations tested showed slightly different levels of resistance and susceptibility in ScCD and CnCD^14^, suggesting that epistasis may have rewired some of the mutation-resistance relationships during the divergence of these two orthologous enzymes.

Here, we set out to fully address the role of epistasis in changing the identity of resistance mutations between ScCD and CnCD. We first report crystal structures of different CD homologs, including distinct conformations of CnCD, and show that all structures are highly conserved in spite of diverging in sequence by up to 60%. We then set out to measure the effects on 5-FC resistance of all single substitutions in both proteins, identify which ones are impacted by epistasis and probe why at the molecular level. We used deep-mutational scanning (DMS) to build comprehensive libraries of single mutants of ScCD and CnCD and expressed them under identical experimental conditions to compare the effect of equivalent mutations in the two orthologs expressed in the same cellular background.

Using machine learning to predict resistance in the pathogen (CnCD) by training the model on DMS data from ScCD, we show that predictions can be made with relatively high-accuracy. By fitting a global epistasis model, we identified a subgroup of epistatic mutants whose within-protein molecular interactions had putatively been rewired in one ortholog but not the other and that are in part responsible for limiting the predictability of the impact of mutations across species. Finally, we characterized the biochemical and structural properties for several pairs of these mutants, allowing us to pinpoint the exact molecular mechanism underlying species-specific resistance.

## Results and discussion

### ScCD and CnCD have highly conserved structures

In order to investigate differences we could expect from mutational effects in ScCD and CnCD, we first assessed their sequence and structural conservation. We have previously shown that the structure of ScCD is very similar to the AlphaFold 2^22^ predicted structure of CnCD^14^ as they share 39% sequence identity. Since then, we reported additional structures of ScCD, including its open and closed conformations, which were key for interpreting the impact of single and pairs of mutations on activity^15^. Using similar approaches, we crystallized and solved the structure of 3 CD orthologs from clinically relevant fungal pathogens, namely *C. neoformans* (CnCD), *Aspergillus niger* (AnCD) and *Nakaseomyces glabratus* (NgCD), at resolutions of 1.50 Å, 1.37 Å and 1.94 Å, respectively. We attempted to work with the *A. fumigatus* (AmCD) ortholog but contrary to AnCD, it could not be purified when expressed in *E. coli*. We thus proceeded with AnCD as a close ortholog and representative of the *Aspergillus* species complex. Together with ScCD, the 4 orthologs have highly conserved structures (Fig. 1c and Supplementary Fig. 1).

CnCD is the only ortholog for which we repeatedly observed mixed conformational states, with one subunit in a closed conformation and the other in an open one (Fig. 1d), similarly to one of our recently reported structures of ScCD (PDB 8VLK)^15^. In contrast, only closed-closed conformations were obtained for AnCD and NgCD. This may reflect differences in conformational dynamics, including transition kinetics, under crystallization conditions. We compared the crystal structures of CnCD (165 residues), AnCD (148 residues), and NgCD (155 residues) with AlphaFold 3^23^ predicted models, and observed closed agreement, with r.m.s.d. values of 0.29 Å over 143 Cα atoms for AnCD, 0.97 Å over 145 Cα for NgCD, and 0.46 Å over 161 Cα atoms and 0.47 Å over 151 Cα atoms for the closed and open conformations of CnCD, respectively. Most differences were mainly localized to the N- and C-termini. AlphaFold 3 predicted with high accuracy the closed conformations for all 3 orthologs. The experimentally observed open state of CnCD was not represented in the prediction, suggesting that functionally relevant C-terminal dynamics could not be captured by AlphaFold 3.

Superposition of the two subunits from our CnCD crystal structure shows a r.m.s.d. of 0.46 Å across 149 common Cα atoms. Structural differences are primarily localized to the C-terminal α-helix, which is displaced from the dimer interface in the open subunit and showed increased flexibility, with terminal residues being disordered. Structural alignment of the dimers of ScCD and CnCD reveals a highly conserved overall fold (r.m.s.d. = 1.01 Å over 278 Cα atoms), including nearly identical configurations for the active site (Fig. 1d). The largest deviations were observed at termini, with the N-terminal region adopting different conformations in each structure and the C-terminal region of CnCD appearing more open and disordered in the open state compared with ScCD (Fig. 1d). This difference in C-terminal regions may, at least in part, arise from the four additional residues in CnCD.

### Most amino acid substitutions have the same phenotypic consequences in ScCD and CnCD

Next, we used DMS to measure the impact of all single amino acid substitutions in ScCD and CnCD. As CnCD perfectly complements the loss of ScCD in *S. cerevisiae*^14^, the impact of mutations on protein function can be directly measured by expressing them in this model yeast. We generated mutant libraries by integrating DNA fragments directly into the yeast genome at the *FCY1* locus by CRISPR-Cas9 genome editing (Extended Data Fig. 1), preserving the endogenous transcriptional regulation of the gene. The mutant libraries were then screened in parallel, using a 5-FC concentration that inhibits around 50% of wild-type growth (Supplementary Fig. 2). Cells were collected at several time points: once before drug exposure (T0) and three more times over the course of the screen. Using deep-sequencing, we converted the change in allele frequencies between T0 and T2 (i.e. after around 10 doublings) into selection coefficients, a metric informing on the fitness effect of a particular mutant relative to the wild-type, with positive effects reflecting higher fitness or 5-FC resistance.

Our experiment was highly replicated and reproducible (Spearman correlation coefficients higher than 0.87 for all pairwise replicate comparisons, Supplementary Fig. 3). First, both libraries were constructed independently twice, in a *MATa* strain and in a *MATɑ* strain. Second, most amino acid variants were represented by two or more codons within each library. Since the genes were split into three fragments, a requirement for high-throughput sequencing, mutants in the fragment overlaps provided additional data points to assess reproducibility. We observed high reproducibility at each level of replication.

In order to balance biological interpretability of our fitness effects estimations with robustness across experimental conditions, selection coefficients were scaled between 0 and 1, using the wildtype and nonsense mutants as reference, respectively. As a result, a resistance score close to 0 indicates 5-FC sensitivity as conferred by the WT enzymes, while a resistance score close to 1 indicates 5-FC resistance, as conferred by complete gene loss. We found that the distribution of fitness effects was strongly bimodal (Supplementary Fig. 4a), corroborating our previous studies in which the same concentration of 5-FC was sufficient to identify resistant mutants^14,15^.

In general, we found a strong correlation in the mutational effects when comparing scores in the overlaps from two fragments (Spearman correlation coefficients upward of 0.83), as well as between the two mating types (Spearman correlation coefficients of 0.87 and 0.88 for ScCD and CnCD respectively, Supplementary Fig. 4b). In downstream analyses, we proceeded differently depending on the parameter studied. To study 5-FC resistance, we first averaged between the two mating types, then classified the mutants as either resistant or sensitive based on a z-score threshold, and finally we calculated the difference between CnCD and ScCD to assess which mutants behaved differently depending on the species. To study epistasis, we instead considered the two mating types as individual replicates, by first confirming that the differences in scores correlated well between mating types (Supplementary Fig. 4c).

Our DMS data for ScCD correlated very well with previous results obtained using the same libraries screened in a different experiment (Spearman correlation coefficient of 0.81, Supplementary Fig. 5a)^14^. This time, 1,350/3,127 (43%) mutants were classified as resistant to 5-FC (Fig. 2a), compared to 32% reported previously^14^. This difference was caused by a lower threshold on resistance scores used here that could be afforded due to the higher replication and higher sequencing coverage and thus better resolution. If we exclude substitutions introducing the wild-type residue of either species and substitutions for which the equivalent was missing in the other species, 1,337/2,998 (45%) mutants were classified as resistant (Extended Data Fig. 2a).

**Figure 2.**
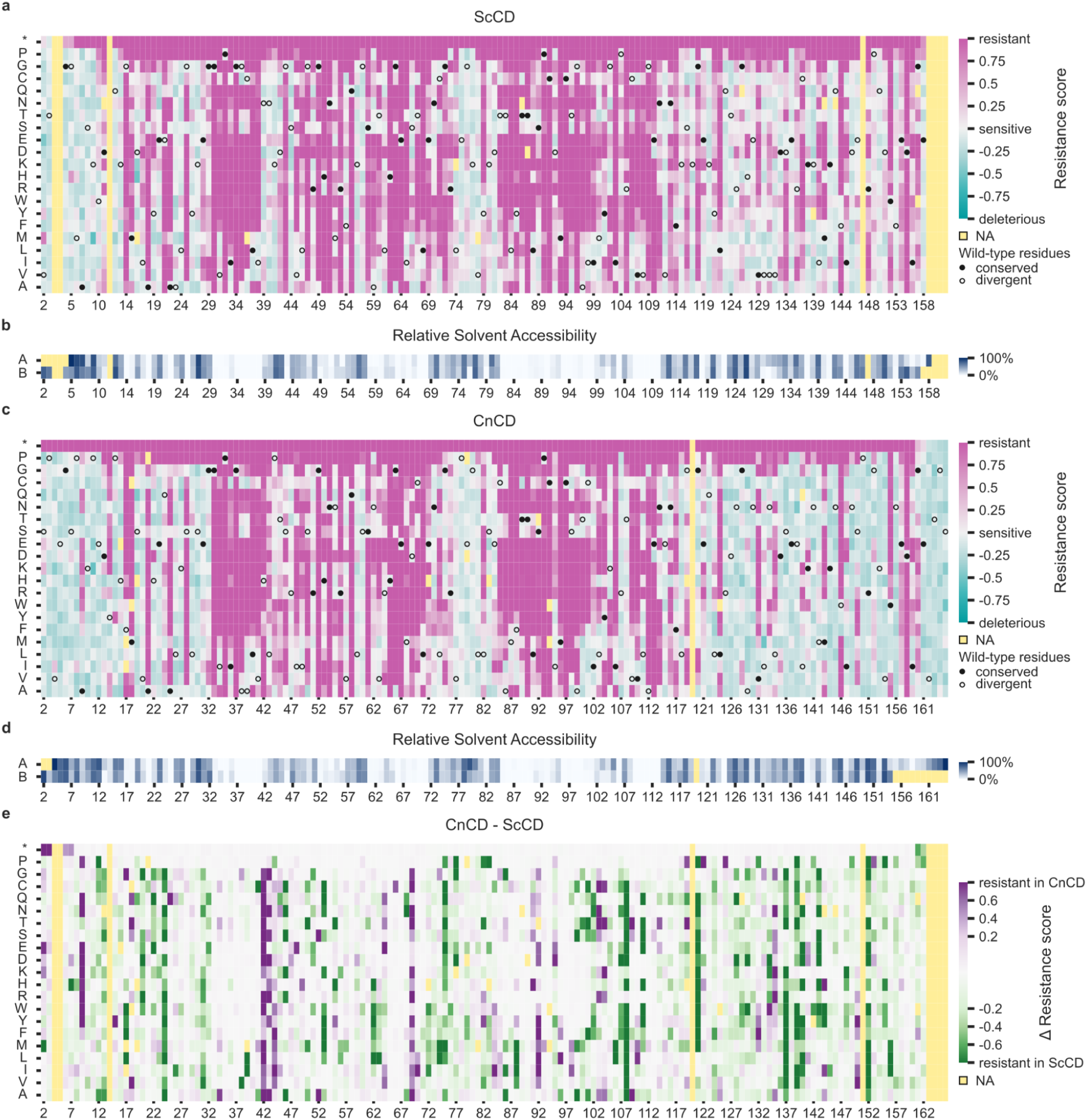
Comprehensive landscape of mutational effects on two fungal cytosine deaminases (CD). Resistance to 5-FC (median of synonymous codons per amino acid) of individual mutants in ScCD (**a**) and CnCD (**c**). The selected loss-of-function (resistant) mutants are colored pink. The wild-type sequences are indicated by circles (filled for conserved sites). Some mutants have scored higher than the manually set upper bound of 1 but were colored as if their score was 1. (**b**, **d**) The relative solvent accessibility is indicated for each chain (A or B) of each protein. In panels a-d, positions indicated on the x axis refer to the corresponding protein sequence. (**e**) Difference in resistance scores (same data as panels a and c), with similarly set bounds on the color scales (-0.8 to 0.8). Purple indicates that mutations introduced in CnCD lead to a LOF (5-FC resistant), while they are functional in ScCD (5-FC sensitive). Green is the opposite, while mutants colored white have the same effects in both enzymes (both are either resistant or sensitive to 5-FC). In this panel, the x axis corresponds to positions in the protein sequence alignment. In all panels, missing data indicated by a pale yellow square can indicate a gap in the alignment (complete column is colored). In panels a, c and e, it may indicate that the mutants did not meet the read count threshold in all T0 replicates. In panels b and d, it may indicate that the position could not be included in the solved structure.

The mutational landscapes for ScCD and CnCD largely overlap (Fig. 2a-e). Among the CnCD mutants,1,323/2,998 (44%) were classified as resistant, with 1,123 of them (85%) being also resistant in ScCD (Extended Data Fig. 2a). This is consistent with the few mutations investigated previously, whose maximum growth rate in 5-FC also correlated very well with our resistance scores (Spearman correlation coefficient of 0.92 and 0.94 for ScCD and CnCD, respectively, Supplementary Fig. 5b).

There was a strong correlation in the impact of single amino acid substitutions on ScCD/CnCD, with a Spearman correlation coefficient of 0.78 when comparing substitutions at sites that are conserved (same amino acid at that position in both wild-type sequences) and 0.66 at sites that are not (Extended Data Fig. 2b). This is expected since conserved sites are less likely to have undergone significant rewiring of their network of intramolecular interactions than those that have been substituted since the split from the common ancestor of the two species. The absolute difference in resistance scores can also be compared at conserved and divergent sites. Expectedly, we found a small but highly significant smaller absolute difference for conserved sites than for divergent sites (Extended Data Fig. 2c) (Mann-Whitney r=0.160, p-value=2.3e-18).

Differences made evident by simply plotting the resistance scores include nonsense mutants before position 6 conferring resistance in CnCD but not ScCD, due to the presence of an alternative start codon in the latter. Additionally, none of the substitutions of the four additional residues in the C-terminal region of CnCD conferred resistance. In most analyses, we have excluded substitutions introducing the wild-type residue from either species to prevent redundancy in the dataset. In general, we expect a low frequency of substitutions to the homologous wild-type residue that would cause loss of function. However, we identified ten such cases in our dataset (G14F, G43S, C71L and G96A conferring resistance in ScCD; F17G, H42N, L43N, L82Y, I95D and G152P conferring resistance in CnCD), with G14F/F17G constituting a reciprocal pair. This indicates a clear rewiring of key interactions involving these residues, providing insight into the factors constraining CD evolution. At the other end of the spectrum, some divergent sites show drastically different patterns between species. For example, most substitutions of the glycine at position 118 in ScCD confer 5-FC resistance, whereas few of them do at the homologous position in CnCD (120, Fig. 2). Similarly, half the substitutions of the arginine at position 148 in ScCD confer 5-FC resistance, while none of them do at the homologous position in *C. neoformans* (R151).

### The resistance phenotype in CnCD can be accurately predicted from ScCD mutational data

We next examined whether the resistance phenotypes in CnCD could be accurately predicted using protein features and training a model using mutational and phenotypic data from ScCD. Such an approach would drastically reduce the efforts required to estimate resistance from any mutation in a pathogenic organism, i.e., an organism often lacking crucial genetic editing tools to quickly test mutational effects in the lab. We computed several features (Supplementary Table 1) directly from the protein structures solved experimentally (e.g. relative solvent accessibility), some from the patterns of evolution of orthologs across species (e.g. GEMME scores) and finally, features that predict the putative impact of substitutions on protein function (e.g. predicted folding energy) including scores from variant effect predictors such as ESM^24^.

We trained a model on the DMS data of ScCD using these features to predict 5-FC resistance. Mutant selection coefficients from the DMS experiment were binarized into resistant or sensitive classes. For each species analyzed separately, the resistance threshold was defined as two standard deviations below the mean score of nonsense mutants. Mutants with scores above this threshold were classified as resistant, whereas those below were considered sensitive. The dataset was split into training (75%) and test (25%) sets, resulting in a classification accuracy of 90%. The most contributing features were the computationally predicted effect of mutations on stability^25^ and ESM scores^24^. We then used the trained model to predict resistance mutations in CnCD, using the DMS data of CnCD to assess the accuracy of the predictions. In general, the model performed well, with 87% accuracy (Extended Data Fig. 3). We also performed the reciprocal analysis, training a model on CnCD, which reached 91% accuracy on the held-out CnCD test set. When applying this model to predict mutation phenotypes in ScCD, it achieved 87% accuracy, indicating that predictive features generalize across species (Supplementary Fig. 6).

Data acquired on one ortholog are therefore capable of being transferred to another one following 600 My of divergence but significant discrepancies remained. A fraction of those could be caused by epistasis, which would prevent extrapolating from one ortholog to the other. Indeed, when predicting CnCD phenotypes from a model trained on ScCD, while the phenotypes of 891/1074 (83%) mutants that were resistant in both orthologs were correctly predicted, the phenotypes of 128/214 (60%) mutations that had opposite effects in the two orthologs were correctly predicted. Some of the opposite in the phenotypic outcomes of mutations between the two orthologs can thus be predicted from these features, but not all. Importantly, the variants that were misclassified generally had weaker phenotypic effects (Supplementary Fig. 7), which means that ambiguous classifications due to weaker effects of mutations could contribute to these wrong classifications.

We also examined which features were significant for correctly predicting the mutations that due to epistasis, changed phenotype between species. For instance, one substitution would be predicted as causing resistance in ScCD but would be predicted as being susceptible in CnCD, presumably because some features have significantly changed between the two orthologs, changing the classification. To investigate this, we analyzed SHAP values for the correctly predicted epistatic mutants. The same core features that dominated the global model—such as ddG stability, ESM and GEMME scores, and residue solvent accessibility—also contributed most strongly to these classifications, and they exhibited similar directional effects. This indicates that the model relies on general biophysical and evolutionary trends. As a result, while these features support the prediction of some phenotype-switching mutations, other cases may require information not captured by the current feature set.

We correctly predict 50% of variants for which substitution to an amino acid that is the wild-type residue at the homologous position results in loss of function. Inspection of the most informative features using SHAP values for this subset of variants reveals that correct predictions are primarily driven by the *ΔΔG* feature estimated using FoldX, which captures the destabilizing effects of these mutations arising from differences in the local structural environment. In contrast, features based on evolutionary conservation, such as GEMME and ESM scores, are misleading in these cases, as they incorrectly favor the homologous wild-type residue as being susceptible and thereby bias predictions toward sensitivity (Supplementary Fig. 8).

As a resource for the fungal antimicrobial resistance community, we predicted the resistance phenotypes for all possible mutants in seven additional CD orthologs from fungal pathogens in which at least one mutation leading to antifungal resistance have previously been observed (Supplementary. Table 2). To account for phylogenetic proximity, we applied the model trained on the CnCD data to predict resistance in the *Cryptococcus gattii* ortholog. Similarly, the model trained on the ScCD data was used to generate predictions for *Nakaseomyces glabratus*, *Candida albicans*, *Candida dubliniensis*, *Candida lusitaniae*, *Candida auris* and *Aspergillus niger*. Interestingly, from the eight single amino acid substitutions associated with resistance phenotypes in the FungAMR database for these orthologs, all were correctly predicted as resistant by the models^7^. We rely on AlphaFold structures for these predictions, but as shown above, these structures are highly accurate, at least for the closed conformation of the enzyme. The complete set of predictions is provided in Supplementary Table 3.

### Contribution of global epistasis to ortholog-specific response to mutations

In order to first get an estimate of the impact of epistasis on mutation-resistance relationships, independently of the predictions, we subtracted resistance scores of equivalent mutants in ScCD and CnCD (Fig. 2e). Again, the values correlated very well with the difference in maximum growth rates between mutants of the two orthologs obtained from previous small scale growth assays^14^ (Spearman ⍴=0.78, p-value=2.0e-5, Supplementary Fig. 5c). Overall, we noted a bias toward slightly more substitutions leading to resistance in ScCD (Fig. 2e), suggesting a global effect on mutation sensitivity or global epistasis. In this case, mutations could have the same impact on the proteins themselves, but the magnitude of the effect would change when measured at the organismal level (or upon heterologous expression) because of the non-linear relationship between growth and enzymatic activity or due to slight differences in experimental conditions.

We considered several mechanisms to explain this difference. One possibility would be that ScCD and CnCD differ in their abundance when expressed in *S. cerevisiae* such that, for the same destabilization, ScCD would reach a lower protein abundance and thus be more likely to confer a resistance phenotype. However, by tagging the two orthologs and measuring their protein abundance in vivo, we found that ScCD was slightly more abundant than CnCD (Supplementary Fig. 9), thereby ruling out this mechanism. We then hypothesized that the proteins might also differ in their stability, such that a mutation with the same destabilizing effect would result in a larger decrease in protein abundance for one of the orthologs. To test this possibility, we expressed both orthologs in *E. coli* and purified the proteins (Supplementary Fig. 10) to measure their melting temperature. We found that the two orthologous proteins have slightly different melting temperatures (T_m_; ScCD, 45°C; CnCD, 43°C)(Supplementary Fig. 11a). Again, a lower melting temperature for CnCD, which we assume translates to lower folding stability, would suggest that CnCD would be more sensitive to mutations, i.e. the opposite of the bias observed in our DMS. Finally, we investigated whether the two enzymes could differ in their inherent biochemical properties, which would result in one causing higher intrinsic sensitivity to 5-FC than the other. We assessed in vitro activity by monitoring substrate disappearance. Under identical conditions, CnCD exhibited reduced apparent activity compared to ScCD (Supplementary Fig. 11b). However, their dose-response curves being very similar indicates that both enzymes behaved the same way in vivo or that their difference in activity results in similar phenotypes (Supplementary Fig. 2). We did note that the IC50 for both orthologs was higher in one mating type, although this may simply be due to strains being selected with two different auxotrophic markers resulting in slightly different metabolisms. Further work will be required to investigate why ScCD appears more sensitive to 5-FC than CnCD in the DMS, seemingly contradicting observations that CnCD is both less stable and less abundant than ScCD.

### One in ten mutations introduced in ScCD and CnCD are epistatic

We sought to distinguish mutants displaying specific epistasis to examine the molecular bases of the changes in phenotypes as specific epistasis indicates which mutants behave differently in ScCD and CnCD because of rewired intramolecular interactions. We hereafter qualify individual mutations as being “epistatic” if they ultimately have background-dependent effects that have been shown not to result from global epistasis or other experimental effects^1^. In order to identify such mutants, we needed to distinguish effects differing between species because of general factors such as experimental noise, from a biologically relevant shift in function of a given amino acid at a given position. This is referred to as a shift in latent phenotype, since we do not know exactly what causes a change in behavior from one species to another.

To infer this shift in latent phenotype from the DMS data, we used a global epistasis model^26^, considering the mating types as individual biological replicates. Once we calculated these values for all missense mutants, we identified 223/2,916 (8%) for which the resistance score significantly differed from one ortholog to the other (adj. *p*-value < 0.001, Benjamini-Hochberg correction, empirical test, Fig. 3a). Since we inferred these differences after modeling the nonlinearities in the data, these shifts in resistance scores represent cases of specific epistasis that are controlled for the differences caused by global epistasis. This single value for the proportion of epistatic mutants is a conservative estimate as different parameter values for the global epistasis model, as well as chance involved in the classification (based on bootstrapping) typically led to proportions ranging from 8 to 12% (approximately one in ten mutations). This is comparable to the proportion of mutants misclassified as resistant or sensitive to 5-FC by our machine-learning model, among which 85 of 349 (24.4%) were epistatic, compared with 123 of 2255 (5.5%) correctly classified variants (Supplementary Fig. 7ab), further showing species-dependent effects are not trivial to identify and are likely caused by a complex combination of factors.

**Fig. 3.**
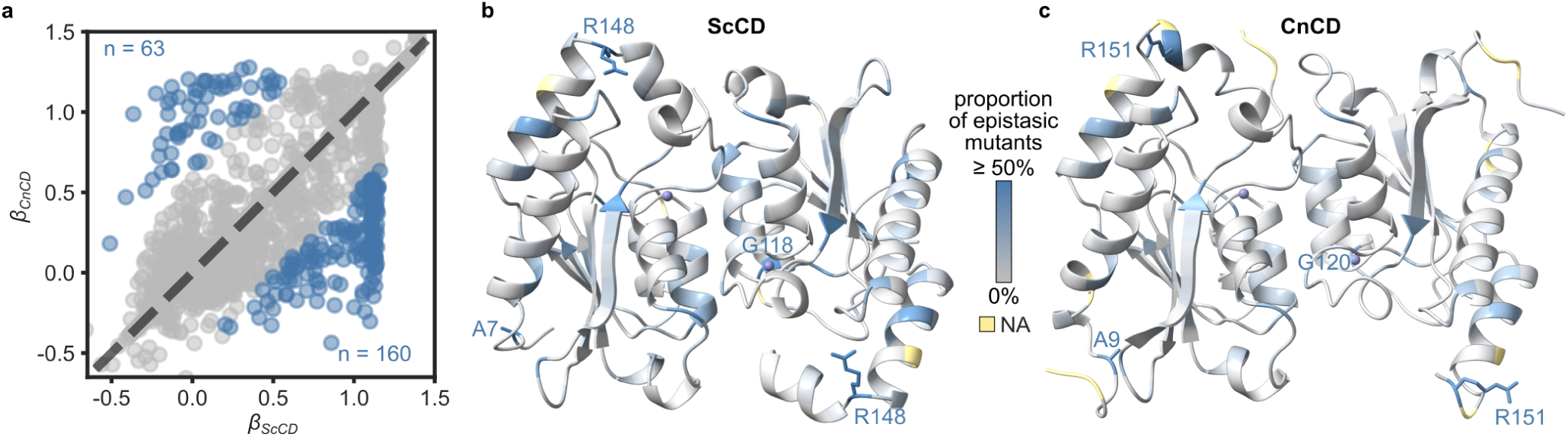
Epistasis landscape of ScCD and CnCD. (**a**) Scatter plot showing the correlation between mutational effects in ScCD and CnCD, inferred by fitting a global epistasis model on the DMS data. Significant epistatic mutants (adjusted p-value below 0.001, see methods section) are highlighted in blue. (**b**, **c**) Proportion of epistatic mutants at each position mapped onto the structure of (**b**) ScCD (PDB 8VLK) and (**c**) CnCD (PDB 36FA). The atoms of residues for which more than half of the substitutions at that position were classified as epistatic are shown, with an accompanying label. Figure generated with ChimeraX v1.8.

Protein regions where we expect epistasis include regions close or directly involved in protein function (e.g. the active site), as seen for other enzymes ^27^, or protein stability (e.g. at the dimer interface). Alternatively, these sites could have been under stronger purifying selection, thereby leaving fewer possibilities for biochemical changes to accumulate and modify the impact of the mutations we introduced. The 223 mutants classified as epistatic cover 83 distinct positions in the protein sequence. However, few key sites, all conserved between ScCD and CnCD, show more than 50% epistatic mutants, namely A7, G118 and R148 in ScCD (A9, G120 and R151 in CnCD) (Fig. 3b,c), some of which we will dissect in more detail below. G118 is located in the active site, but the two other residues reside at the periphery of the dimer, far from the active site or the dimer interface, with no obvious link with the active site or protein stability as these residues are not buried. The dimer interface does not appear to be enriched for epistatic positions either as the fraction of interface positions harboring at least one epistatic mutation is similar to that of non-interface positions (35/65 vs 48/90 respectively; χ² test, p ≈ 1). Together these results show how the rewiring of genotype-phenotype relationships can happen at numerous sites in the protein.

### The molecular bases of epistasis in CD

We hypothesized that local structural context of substitutions drives their phenotypic effects and epistasis. We selected representative mutants for biophysical and biochemical characterization from conserved ScCD/CnCD sites with high epistasis (>50%) and from variants spanning resistance phenotypes and structural locations, including five of seven variants misclassified by the machine learning model. We set out to purify and assay the function of several mutants in the two orthologs to validate the impact of mutations in vitro and attempt to rationalise the bases of epistasis. Pairwise comparisons were performed for six equivalent mutants in ScCD and CnCD, where each pair comprised one variant associated with a resistance phenotype in one ortholog and the corresponding variant associated with susceptibility in the other, with mutations distributed across the structure (Fig. 4). Across these comparisons, thermal denaturation assays consistently supported the hypothesis that resistance-associated substitutions destabilize protein structure (Fig. 4). In five cases out of six, resistance-associated variants exhibited severely reduced yields of soluble protein following heterologous expression in *E. coli* relative to their corresponding wild-type proteins (Supplementary Fig. 10), suggesting defects in folding or reduced stability. Stability measurements led to the absence of detection of T_m_ for these five selected mutants, indicating that the majority of these proteins were unfolded (Fig. 4 and Supplementary Fig. 12). Overall, these results support a model in which resistance is frequently associated with strong structural destabilization in vitro, and the ortholog-specificity of these phenotypes in vitro reflects the resistance phenotypes. The last case, inserting a serine at the homologous positions 99 and 96 led to proteins with stabilities near WT levels but ScCD did not have detectable activity, while CnCD retained minimal activity, which appears to be sufficient to confer susceptibility to 5-FC in vivo. This is consistent with recent findings showing that slightly higher than 5% of activity in ScCD is sufficient for 5-FC to be toxic^28^.

**Fig. 4.**
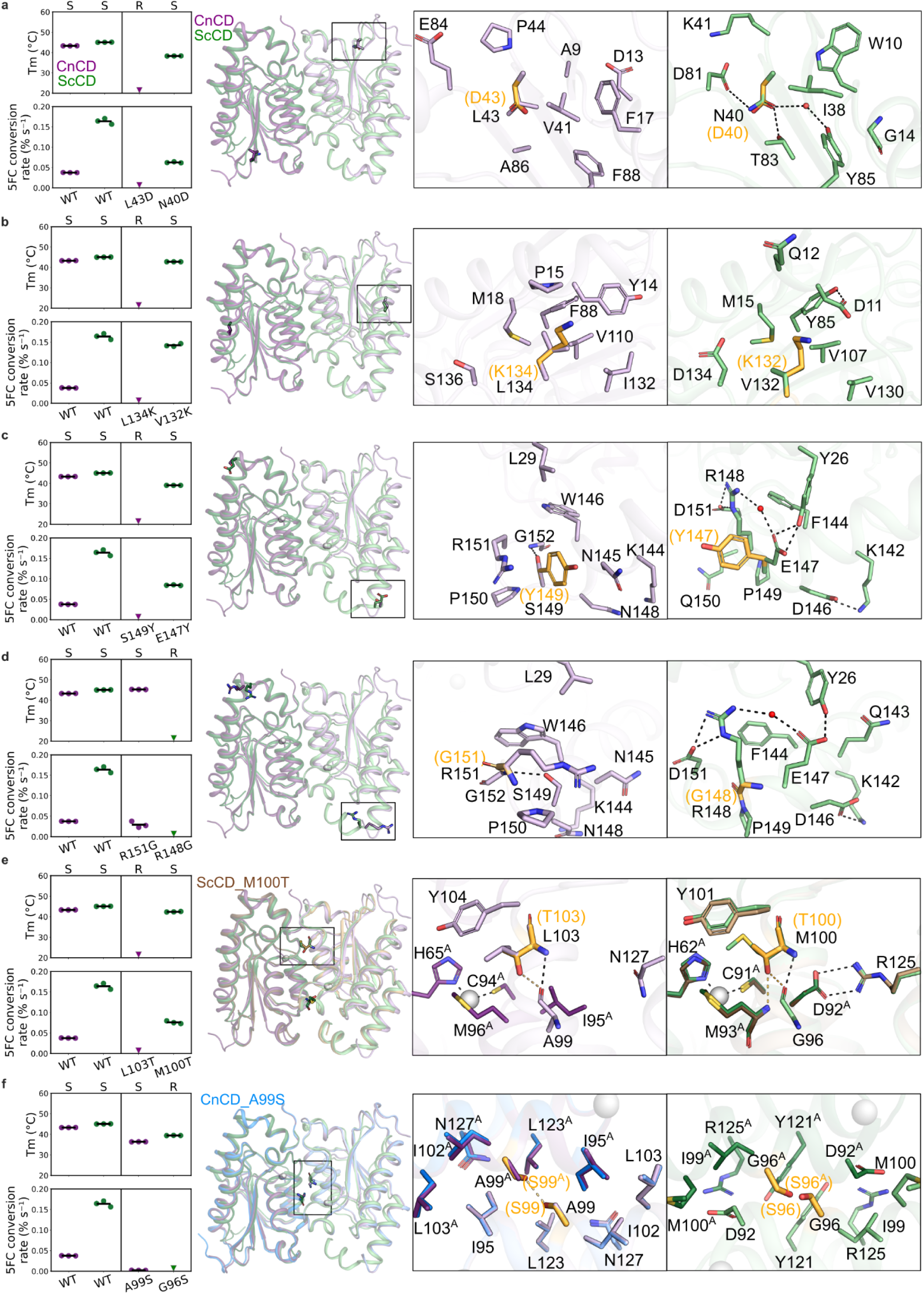
Thermal stability, activity, and structural context of CnCD and ScCD variants. (**a-f**) *Left*: Top, melting temperature (T_m_) measured by nanoDSF. 5-FC-sensitive (S) or -resistant (R) phenotypes from DMS are shown above. Bottom, activity expressed as the slope of 5-FC conversion over time. Variants are grouped in pairs corresponding to equivalent mutations in CnCD (purple) and ScCD (green). Triangles indicate variants with no detectable unfolding transition or no detectable activity. Each point represents an independent measurement (n = 3); horizontal bars indicate the mean. *Middle*: Overall structures of wild-types CnCD (purple, PDB 36FA) and ScCD (green, PDB 8VLK, subunits A and B) shown as cartoons, with experimentally determined mutant crystal structures shown for ScCD_M100T (brown, PDB 36GM) and CnCD_A99S (blue, PDB 36FG). Residues of interest are shown as sticks within the boxed region. *Right*: Local environments of each substitution in CnCD and ScCD, with substituted side chains overlaid onto the wild-type structures. Carbon atoms of substituted residues are shown in light orange (orange labels and brackets). Zinc and water are shown as gray and red spheres, respectively. Dashed lines indicate polar interaction networks. Unless marked with a superscript A, stick residues belong to subunit B. (a) CnCD_L43D/ScCD_N40D. (b) CnCD_L134K/ScCD_V132K. (c) CnCD_S149Y/ScCD_E147Y. (d) CnCD_R151G/ScCD_R148G. (e) CnCD_L103T/ScCD_M100T. (f) CnCD_A99S/ScCD_G96S. Protein structure panels were generated using PyMOL v3.1.0^21^.

Examining the location of these residues allows us to rationalize why these mutations have ortholog-specific effects. The pair CnCD_L43D (resistant) and ScCD_N40D (sensitive) illustrates how the local environment can influence whether the same physicochemical change brought by mutation is tolerated or not. The CnCD_L43D variant yielded around four times less soluble protein than the wild-type (Supplementary Fig. 10), whereas the equivalent ScCD_N40D had similar yield to ScCD_WT and retained partial activity despite reduced thermal stability (ΔT_m_ ≈ −7 °C). Structural analysis provides a likely explanation: in CnCD, L43 is embedded in a tightly packed hydrophobic cluster formed by the side chains of A9, F17, V41, P44, A86 (Fig. 4a). Substitution to aspartate introduces an unfavorable charge into this compact, non-polar environment, disrupting local packing and compromising structural integrity. In contrast, ScCD N40 lies in a more polar environment and participates in a water-mediated hydrogen-bonding network with K41, D81, T83, and Y85 (Fig. 4a). Here, substitution to an aspartate preserves the polar character of this region, limiting structural disruption, although loss of the asparagine-specific hydrogen bond reduces stability. Consistently, DMS data shows strong intolerance to polar or charged substitutions at this position in CnCD, whereas similar substitutions are broadly tolerated in ScCD (Fig. 2a, 2b).

A similar context-dependent effect is observed for CnCD_L134K (resistant) and ScCD_V132K (sensitive). In CnCD, L134 is located within a constrained hydrophobic environment shaped by residues from the first α-helix, with P15 introducing a bend that brings this region into close proximity (Fig. 4b). Substitution to a lysine is destabilizing, likely because the restricted environment cannot accommodate a charged side chain without disrupting local interactions. Consistently, CnCD_L134K yielded only trace amounts of soluble protein (Supplementary Fig. 10) and showed no detectable T_m_, indicating severe destabilization. In contrast, the equivalent position in ScCD is less constrained, and likely allows partial solvent exposure of the lysine, consistent with a susceptible phenotype (Fig. 4b).

Another illustrative case involves CnCD_R151G (sensitive) and ScCD_R148G (resistant). In CnCD, R151 is largely solvent-exposed and adopts different conformations in open and closed states, forming a single hydrogen bond with S149 in the open form and none in the closed form (Fig. 4d). Consistent with this limited structural role, DMS data indicate broad tolerance at this position. In vitro, CnCD_R151G has a slight increase in stability (ΔT_m_: +2 °C) and retains near WT activity, suggesting structural permissiveness. In contrast, in ScCD, the equivalent arginine is embedded in a more stable interaction network, including direct and water-mediated hydrogen bonds (Fig. 4d). Disruption of this network by a substitution to glycine, likely leads to the loss of a detectable melting transition and strongly reduced activity (Supplementary Fig. 13). These observations suggest that this residue is structurally important, specifically in this background.

A similar pattern is observed for ScCD_E147Y (sensitive) and CnCD_S149Y (resistant), where differences in local architecture and connectivity again likely influence tolerance to substitutions. In ScCD, E147 is solvent-exposed and interacts with Y26 from a neighboring α-helix (Fig. 4c). E147 also participates in water-mediated contacts with Y26 and R148, and R148, in turn, interacts further with D151 in the C-terminal helix, creating an extended interaction network linking these structural elements. The aliphatic portion of E147 also engages in van der Waals contacts with R148. Substitution of E147 with tyrosine is structurally tolerated, as the tyrosine side chain can remain solvent-exposed, but disruption of the aforementioned key hydrogen-bonding and water-mediated interactions, likely accounts for the observed decrease in stability. In contrast, in CnCD, the corresponding S149 is partially buried between helices and constrained by a local structural arrangement, including a proline residue present in CnCD, but absent in ScCD due to a local indel, which initiates and rigidifies the C-terminal helix. In this context, substitution with tyrosine in CnCD is unfavorable, due to steric incompatibility and inability to reorient toward the solvent without perturbing helix packing, consistent with the absence of a detectable T_m_ and a resistant phenotype in DMS data (Fig. 2e).

Overall, the cases described here suggest that the observed epistasis can be linked to effect on a local subunit environment through either physicochemically different environment (hydrophobic vs polar) or presence and absence of stabilizing interaction networks within a given chain.

Mutations at the dimer interface, in proximity to the active site, further support the idea that the phenotypic consequences of equivalent substitutions depend strongly on local structural context. A clear example is provided by substitutions at the catalytic-site hydrophobic surface. We have solved the crystal structure of ScCD_M100T at 2.97 Å resolution. In ScCD, M100 contributes to the hydrophobic character of the catalytic site, and substitution to threonine is accommodated in this environment: the threonine methyl group occupies a similar position to the methionine side chain, while the hydroxyl group forms stabilizing hydrogen bonds with the backbone carbonyl of G96 (2.7 Å) and the backbone amide nitrogen of M93 (Fig. 4). The D92-R125 salt bridge involved in intersubunit interactions is also maintained in this arrangement. In contrast, in CnCD, the equivalent residue L103 lies within the hydrophobic core of the dimer interface, where substitution to threonine would likely be poorly accommodated due to predicted steric clashes with I95 and the nearby backbone carbonyl of A99 (Fig. 4), while introducing a polar group into a region that normally requires hydrophobicity. These observations further support the notion that resistance-associated substitutions preferentially emerge in structural contexts where equivalent physicochemical changes are poorly accommodated.

Additional evidence for context-dependent effects at the dimer interface comes from a pair of variants in which both orthologs remain folded yet differ functionally: ScCD_G96S shows complete loss of activity, whereas CnCD_A99S retains residual activity (Supplementary Fig. 13). In ScCD, G96 residues from each subunit face one another at the center of the homodimer interface, while surrounding residues participate in a more polar interaction network, including a Y121 self-interaction between subunits, the D92–R125 salt bridge, and D92–I99 van der Waals contacts (Fig. 4e). Substitution to serine would disrupt the symmetric G96-G96 packing, likely introducing solvent access, which would reduce stability (ΔT_m_ = -5.5 °C) and abolish catalytic activity. In CnCD, A99 engages in a similar A99-A99 interface (Fig. 4f). We solved the crystal structure of CnCD_A99S (PDB 36FD) and the structure reveals that the Ser residues are accommodated and the surrounding hydrophobic structural network (such as I95 and L103)(Fig. 4f), likely helps maintain dimer integrity and provides greater tolerance to substitutions to small residues (S, G, or C). Accordingly, A99S reduces stability (ΔT_m_ = -6.9 °C) and activity, but does not abolish function, consistent with serine being accommodated at the interface while narrowing the local conformational space available at this position, which may in turn modulate the propagation of conformational effects through the structure. Together, these observations suggest that subtle perturbations at symmetric interface residues near the active site can differentially propagate to the catalytic center through changes in conformational dynamics.

We also observed a marked divergence in the functional impact of equivalent glycine-to-glutamic acid substitutions in the two orthologs (ScCD_G118E and CnCD_G120E) (Fig. 5a). In ScCD, G118E is associated with a resistant phenotype and exhibits substantial destabilization in vitro, with no measurable T_m_ (Fig. 5a). In contrast, the CnCD_G120E variant exhibited significantly increased stability, with a 10 °C rise in T_m_, accompanied by enhanced catalytic activity (Fig. 5a). We examined crystal structures of both WT orthologs and hypothesized that epistasis arises primarily from differences in how the glutamic acid side chain is accommodated. To test this hypothesis, we solved the structure of the CnCD_G120E variant at 1.73 Å resolution (PDB 36FE, Fig. 5b). The structure confirmed that the E120 side chain is solvent-exposed, where its positioning both shields the hydrophobic patch formed by I95 and L123 of the two subunits from direct solvent exposure and facilitates the buildup of a water network compatible with the protein surface in the surrounding solvent (Fig. 5c). In addition, the aliphatic portion of the glutamate side chain forms favorable van der Waals interactions with I95 and L123, reinforcing local hydrophobic packing (Fig. 5c). In ScCD, however, such an orientation does not seem possible since the glutamate side chain is instead directed toward the helix, likely causing local structural disruption (Fig. 5c). This difference appears explained in part by a single-residue indel immediately upstream of the substitution site. ScCD contains a lysine (K117), whereas CnCD lacks the corresponding residue (Supplementary Fig. 1). Structural superposition shows that, although this region aligns well overall, the lysine in ScCD introduces a kink at the N-terminus of an α-helix, altering side-chain accommodation. Together, these interactions provide a structural explanation for the increased stability and enhanced enzymatic activity observed for CnCD_G120E, while further illustrating how subtle architectural differences between orthologs can profoundly alter mutational outcomes.

**Fig. 5.**
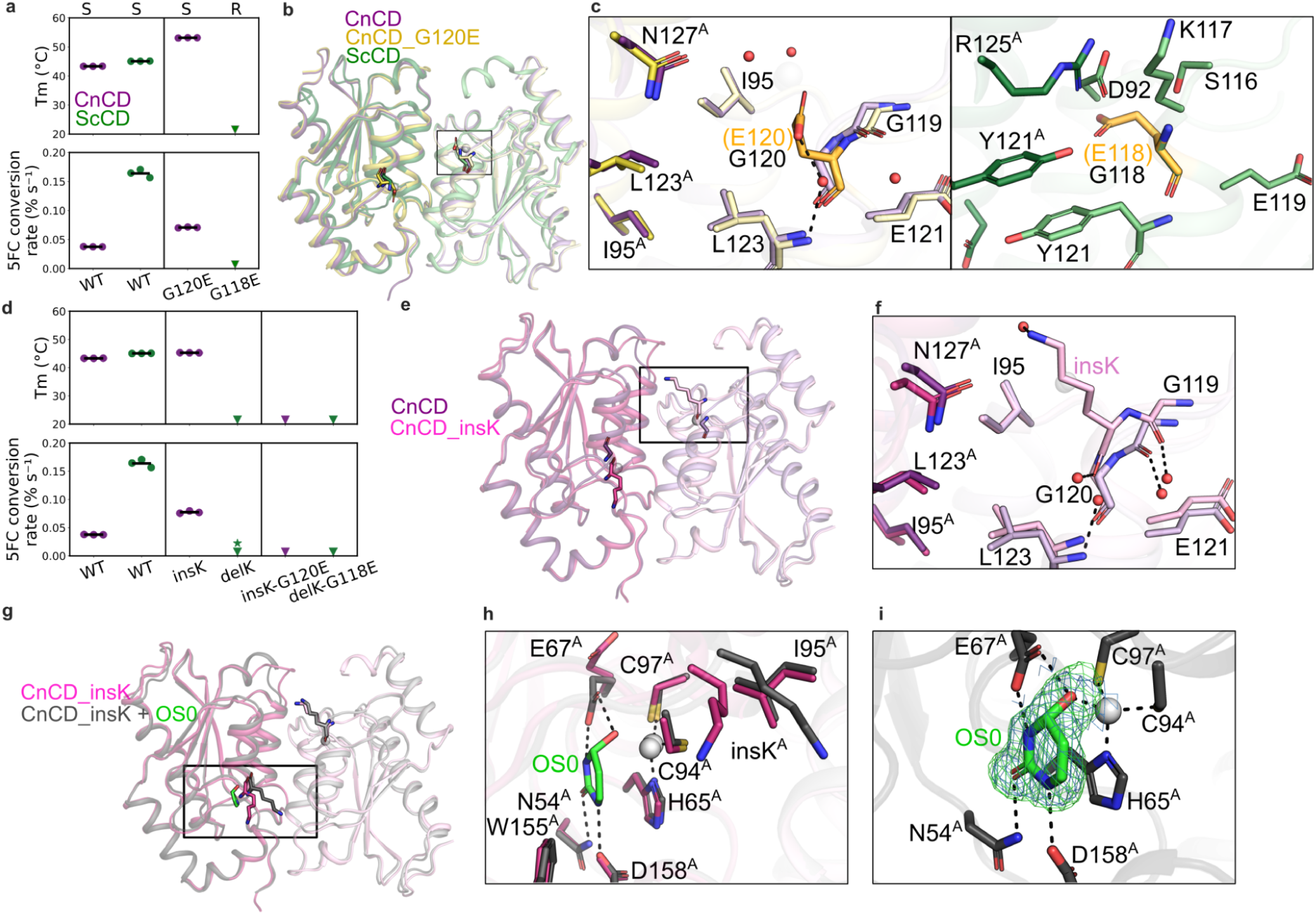
Structural basis of epistatic effects underlying the G118E/G120E substitution. (**a, b**) *Left*: Top, melting temperature (Tm) measured by nanoDSF. 5-FC-sensitive (S) or -resistant (R) phenotypes from DMS are shown above where available. Bottom, activity expressed as the slope of 5-FC conversion over time. Variants are grouped in pairs corresponding to equivalent mutations in CnCD (purple) and ScCD (green). Triangles indicate variants with no detectable unfolding transition. Each point represents an independent measurement (n = 3); horizontal bars indicate the mean**. b**) Overall superposition of crystal structures of wild-type CnCD (purple, PDB 36FA), CnCD_Kins (yellow, PDB 36FF) and ScCD (green, PDB 8VLK, subunits A and B). Residues of interest are shown as sticks within the boxed region. **c**) Local environments of substitutions G120E in CnCD and G118E in ScCD, respectively. Carbon atoms of substituted residues are shown in light orange (orange labels and brackets). Dashed lines indicate hydrogen bonds. Zinc and water are shown as gray and red spheres, respectively. **e**) Overall superposition of crystal structures of CnCD_WT and CnCD_Kins. **f**) Close-up view of the local structural environment of the lysine inserted in CnCD. **g**) Superposition of crystal structures of CnCD_Kins in absence and presence of an inhibitor (CnCD_Kins + OS0). The inhibitor is shown in green sticks. **h**) Local environment of inhibitor OS0 in CnCD_Kins. **i**) Close-up view of the electron density around OS0. The 2Fo–Fc map is shown as a blue mesh contoured at 1.0σ. The positive Fo–Fc omit map is shown as a green mesh contoured at 2.5σ. All maps are carved at 2 Å around the ligand. Protein structure panels were generated using PyMOL v3.1.0^21^.

As ScCD_G118E appears unable to adopt the productive solvent-exposed conformation observed in CnCD_G120E, likely due in part to the presence of K117, we sought to evaluate directly the contribution of this residue in shaping the difference in the G-to-E substitution accommodation. We engineered reciprocal variants by deleting the K117 in ScCD and introducing the corresponding lysine into CnCD. Deletion of this lysine in ScCD led to no detectable T_m_ and substantial activity loss, indicating that K117 contributes significantly to protein stability in this background. Conversely, insertion of the corresponding lysine into CnCD led to a very modest increase in stability (ΔT_m_ = +1.3 °C) but produced a substantial enhancement in enzymatic activity, suggesting that, while this residue is not essential for structural stability in CnCD, it can positively modulate catalytic performance (Fig. 5d).

Crystal structures of the lysine-inserted CnCD variant, determined in the absence (Fig. 5e-f, PDB 36FF) and presence of 2-hydroxypyrimidine (pyrimidin-2-one), which is enzymatically converted into a tightly bound hydrated active-site adduct that acts as a mechanism-based inhibitor (Fig. 5g-i, PDB 36FG), further support a structural role for this residue. The inserted lysine is associated with reduced solvent exposure of the hydrophobic patch composed of I95 and L123 and a more structured local solvent organization. This resembles the effect of the CnCD_G120E substitution, suggesting partial recapitulation of its stabilizing environment by the indel.

We next introduced the G-to-E substitution into these engineered backgrounds to test whether the indel alone was sufficient to explain epistasis. In contrast to the pronounced gain in stability and activity observed for CnCD_G120E, the same substitution in the lysine-inserted background (CnCD_insKG120E) resulted in no detectable T_m_ or catalytic activity (Fig. 5d). Similarly, the ScCD_KdelG118E variant also exhibited no detectable T_m_ and no measurable activity, indicating that removal of K117 is insufficient to enable productive accommodation of the glutamate substitution in ScCD. Together, these results demonstrate that although the presence or absence of this lysine contributes to the differential accommodation of the glutamate side chain, the indel alone is insufficient to fully recreate the ortholog-specific mutational effects. Instead, the broader local structural environment appears to determine whether this resistance-associated substitution can be productively accommodated.

## Conclusion

Because of epistasis, amino acid substitutions in orthologous proteins can have dramatically different outcomes ^9,29^. A spectacular outcome are compensated pathogenic deviations, which are pathogenic amino acid substitutions that correspond to wild-type residues in orthologous proteins^3,30^. One particular context in which this is important is for drug resistance phenotypes, whereby multiple species are often treated with the same drugs. Because of epistasis, different sets of mutations could cause resistance in orthologous proteins, even if the mechanisms of resistance are the same. In fungi, some mutations in the targets Erg11 and Fks1 have been reported to respectively cause resistance to azoles and echinocandins in multiple fungal species^7^, suggesting that the identity of resistance mutations could be conserved across millions of years of evolution. However, no data allows to fully assess the extent of conservation as data is too sparse and there is publication bias towards reporting resistance mutations. Here, using the fungal CD as a model, we examine whether drug resistance mutations are conserved between two orthologous proteins by measuring the impact of all possible mutations on resistance.

The homodimeric enzyme CD is responsible for the deamination of the prodrug antifungal 5-FC (flucytosine), converting this pro-drug into a toxic one. Resistance to this drug can be caused by the nearly complete LOF of the enzyme^7,14^. LOF of CD is sufficient to cause complete resistance to 5-FC in various fungal species^7,14^, although resistance during treatment or experimental evolution can also occur through the LOF of other enzymes in the same metabolic pathway, such as Fur1^11,31^, the enzyme catalyzing the reaction immediately downstream of CD in the pyrimidine salvage pathway.

Despite the 650 millions of years that separate *S. cerevisiae* and *C. neoformans*,^32^ resulting in 61% protein divergence between the two orthologs, LOF of CD and therefore 5-FC resistance, appears attainable mainly by the same amino acid substitutions. This level of conservation in the impact of amino acid substitutions allows one to predict the effect of mutations in one ortholog using a machine learning model trained on the other with accuracy reaching nearly 90%. This enables the interpretation of the impact of mutations in CD across the most important fungal pathogens with high-accuracy, without the need to perform full experimental characterization of mutations. One limiting factor appears to be epistasis. About 10% of substitutions have opposite effects when introduced in the orthologs.

Epistasis can have quantitative and qualitative impact on phenotypes. In the first case, it can change the extent to which a mutation alters protein function. In the second, the same mutation can have completely different qualitative impacts on function, for instance by being neutral or beneficial in one ortholog and pathogenic in the other. An example is the A53T substitution in human ɑ-synuclein, which predisposes to Parkinson’s disease, while mice with a threonine at the homologous site are healthy^3,33^. Epistatic mutations in CD are also qualitative as they are classified as being neutral or adaptive in the presence of 5-FC. This classification is allowed due to the fact that fitness effects are bimodal, most likely resulting from the fact that the relationship between protein activity and fitness is highly steep, requiring nearly a complete loss to confer resistance to 5-FC. Interestingly, our estimates of the fraction of mutations that lead to different qualitative outcomes in CD is close to the estimated rates of compensated pathogenic deviations initially reported in mammals^3^ and insects^30^, suggesting a general trend that is reflective of other proteins.

Using in vitro biochemical assays, we confirmed that epistatic mutations often affect the stability and or activity in an ortholog-specific manner, validating that their impact depends on the sequence of the proteins themselves. Across multiple equivalent substitutions, we show that identical physicochemical changes can lead to distinct biochemical outcomes, ranging from severe destabilization to near-wild-type behavior, depending on the local structural environment of the protein in which they occur. In many cases, resistance-associated substitutions are linked to pronounced destabilization or loss of folding, resulting from the disruption of local interaction networks, while the same changes are tolerated in alternative structural contexts that provide compensatory interactions or reduced steric constraints.

Importantly, our results further demonstrate that epistasis is not solely determined by single-residue differences but is strongly influenced by local architectural features of the protein. In particular, we show that indels can remodel secondary structure geometry and alter side-chain accommodation, thereby reshaping the local environment in which mutations are interpreted. This remodeling can modulate the extent to which equivalent substitutions are stabilized, tolerated, or destabilizing, and in some cases enables or prevents the formation of compensatory interaction networks such as water-mediated hydrogen bonding or hydrophobic packing rearrangements. However, our reciprocal engineering experiments also indicate that indels alone are insufficient to fully account for the divergence in mutational outcomes, implying that epistasis arises from a distributed set of structural determinants rather than a single localized feature.

The phenotype we examined here is nearly binary, owing to the fact that the cytosine deaminase has a relationship between function (approximated by dosage) and resistance that has a steep threshold and that resistance derives from LOF^28^. Knowing whether our findings apply directly to other resistance mechanisms will await further investigation as this relationship has not been characterized for any other resistance proteins. For most genes involved in antifungal resistance, we know very little about the quantitative relationship between activity and resistance. The fact that LOF in CD is required for resistance makes the finding not applicable to the important antifungal drug targets Erg11 and Fks, as these need to remain functional for resistance. However, LOF leading to resistance is a common mechanism outside of CD. For instance, resistance to polyenes can emerge from LOF mutations in multiple enzymes in model yeast and pathogens ^34,35^. In a recent survey of all known antifungal resistance mutations in fungi, we identified stop codons in many target genes (e.g. *ERG3, ERG6, FUR1*) and many resistance amino acid substitutions were predicted to be destabilizing across genes involved in resistance across several classes of antifungals. LOF is thus a frequent mechanism of resistance acquisition, making CD a tractable and representative model and our approach extendable to many other important proteins. Our results show that with data from a single or a handful of species, the impact of mutations could be extrapolated to many other species using machine learning with features that can be derived computationally, alleviating the need for complex experiments.

## Materials and methods

### *E. coli* expression and protein purification

The coding sequences of *FCY1* from *S. cerevisiae*, *C. neoformans, Aspergillus niger, and Nakaseomyces glabratus* were synthesized and subcloned into the pET28a expression vector. Each construct included an N-terminal 6×His tag followed by a thrombin cleavage site, positioned in-frame upstream of the coding region. Site-directed mutagenesis of ScCD and CnCD was carried out using Kapa polymerase (Roche) with the primers listed in Supplementary Table 5. All mutations were confirmed by Sanger sequencing. Plasmids encoding wild-type and mutant variants were introduced into *E. coli* BL21(DE3) chemically competent cells via heat-shock transformation. Individual transformants, or colonies obtained from glycerol stock streaks on LB-kanamycin agar plates when fresh transformants were unavailable, were used to inoculate 10 mL overnight pre-cultures in LB medium containing 50 µg/mL kanamycin. Pre-cultures were subsequently diluted 1:100 into 800 mL Terrific Broth (TB) supplemented with 50 µg/mL kanamycin for large-scale protein expression.

Cultures were incubated at 37 °C with shaking at 200 rpm until reaching an optical density at 600 nm (OD₆₀₀) of 0.6–1.0. Protein expression was induced by the addition of 0.5 mM isopropyl β-D-1-thiogalactopyranoside (IPTG), and 0.5 mM zinc acetate was supplemented at the same time to ensure availability of the zinc cofactor.. Following induction, cultures were shifted to 16 °C and incubated for approximately 18 h with continuous shaking. Cells were collected by centrifugation at 6,000 x *g* for 15 min at 4 °C, and the resulting pellets were stored at −20 °C until further processing.

Cell pellets were resuspended in lysis buffer (50 mM Tris-HCl, pH 7.5; 400 mM NaCl; 2% [v/v] glycerol; 5 mM imidazole) supplemented with lysozyme (0.5 mg/mL), PMSF (0.17 mg/mL), and DNase I (0.01 mg/mL). Cells were lysed by sonication for 10 min at 35% amplitude (10 s on/20 s off cycles), and the lysate was clarified by centrifugation at 21,700 x *g* for 30 min at 4 °C. The supernatant was incubated with cobalt-based IMAC resin for 50 min at 7 °C with gentle agitation. The resin was sequentially washed with a gradient of imidazole. Bound Fcy1 was eluted with 3 mL elution buffer (50 mM Tris-HCl, pH 7.5; 1 M NaCl; 5% glycerol; 200 mM imidazole) following a 10 min incubation. The eluate was buffer-exchanged into PBS (10.1 mM Na_2_HPO_4_,136.9 mM NaCl, 2.7 mM KCl, 1.8 mM KH_2_PO_4_) containing 0.02% Tween-20 using a PD-10 desalting column. For the preparations leading to the CnCD_WT, CnCD_A99S and ScCD_M100T crystal structures, the N-terminal His tag was removed by thrombin digestion (7 U per mg protein). The cleaved proteins, which did not bind Ni-NTA resin, were collected from the flow-through. The final protein samples were either diluted for activity and stability assays or concentrated to 15 mg/mL for crystallization. Protein concentrations were determined by measuring absorbance at 280 nm using a NanoDrop 2000/2000c spectrophotometer (Thermo Fisher Scientific) and calculated using theoretical molar extinction coefficients.

### Protein crystallization, data collection and structure determination

Initial crystallization conditions were screened using commercial kits from NeXtal Biotechnologies (Holland, OH, USA) using the microbatch-under-oil method. Most constructs were crystallized in the presence of His-tag while the His-tag was cleaved via thrombin prior to crystallization setup for CnCD, CnCD_A99S and ScCD_M100T. The detailed reservoir solutions for these crystals are listed below: CnCD (0.01 M zinc chloride, 0.1 M HEPES pH 7.0, 20% (w/v) PEG 6000); AnCD (0.2 M ammonium sulfate, 30 % (w/v) PEG 4000); NgCD (0.1 M MES pH 6.5, 25% (w/v) PEG 6000); CnCD_A99S (0.2 M ammonium chloride, 0.1 M MES pH 6.0, 20% (w/v) PEG 6000); CnCD_G120E (0.2 M lithium chloride, 0.1 M MES pH 6.0, 20% (w/v) PEG 6000); CnCD_Gly119_Gly120insLys (0.1 M MES pH 6.5, 25 % (w/v) PEG 6000); CnCD_Gly119_Gly120insLys complex with OS0 (0.2 M sodium chloride, 0.1 M MES pH 6.0, 20% (w/v) PEG 6000), and ScCD_M100T (0.1 M HEPES pH 7.5, 10% (w/v) PEG 6000, 5% (v/v) MPD). Obtained in the presence of PEG 6000 or PEG 4000, these crystals were cryoprotected either by their reservoir solutions or by the reservoir solutions supplemented with 15% (v/v) ethylene glycol prior to data collection at 100K at the CMCF-ID beamline (Canadian Light Source) or at the LRL-CAT (31-ID) beamline (Advanced Photon Source, Argonne National Laboratory).

The wavelengths for these data collections are shown in Supplementary Table 4. Data processing and scaling were performed with XDS^36^. The structures were determined by MolRep^37^ using the corresponding AlphaFold predicted models or the previously determined structure of ScCD (PDB 8VLM) as the search templates. The models were completed via multiple cycles of refinement using Refmac5^38^ in CCP4^39^ followed by model correction with Coot^40^. The stereochemistry of the final models was analyzed with PROCHECK^41^. Data collection and refinement statistics are shown in Supplementary Table 4. The coordinates and structure factors of CnCD, AnCD, NgCD, CnCD_A99S, CnCD_G120E, CnCD_G119_G120insK, CnCD_G119_G120insK complex with OS0, and ScCD_M100T have been deposited in the Protein Data Bank with the accession codes 36FA, 36FB, 36FC, 36FD, 36FE, 36FF, 36FG, and 36GM, respectively.

### Strain constructions for the DMS experiment

Deep-mutational scanning libraries of ScCD were generated in a previous study^15^. Briefly, the mega-primer method was used to generate codon-by-codon libraries on a high-copy vector bearing the codon-optimized sequence of Sc*FCY1*. After recovery from bacteria, variants were knocked in, position by position, at the genomic locus of *FCY1* in strains BY4741 (*MATa*) and BY4742 (*MATɑ*) of *S. cerevisiae* using CRISPR-Cas9 gene editing. Separate glycerol stocks for each position of ScCD allowed to pool into three overlapping fragments (F1, F2, F3) for amplicon sequencing (F1 = Val 2 to Leu 68, F2 = Val 45 to Lys 115, F3 = Asp 92 to Glu 158).

For CnCD, we first deleted the *ho* locus in BY4741 and BY4742, then deleted *FCY1* by replacing it with a resistance marker (HYG in BY4741 and NAT in BY4742). Cn*FCY1* was divided in 7 fragments of 23 or 24 amino acid residues (F1 = Ser 2 to Gln 24, F2 = Ala 25 to Arg 47, F3 = Ile 48 to Leu 71, F4 = Glu 72 to Cys 94, F5 = Ile 95 to Leu 118, F6 = Gly 119 to Asn 141, F7 = Met 142 to Ser 165). We amplified 7 different gene versions in which each fragment was replaced by a stuffer sequence^42^ using fusion PCR (see Supplementary Table 5 for primers). We used a CRISPR-Cas9 gene-editing protocol targeting NAT or HYG markers to integrate these 7 versions of Cn*FCY1* in the genome of BY4741 or BY4742. Each construction was validated by Sanger sequencing to confirm the proper integration of the construct.

DMS library genomic integration was performed as follows. First, 7 oPools (IDT technologies, one per fragment) were ordered, each containing NNN codons at each position. We used each oPool as a PCR template to generate the mutant library (Supplementary Table 5 for primers). Each oPool was amplified 16 times to help with the uniformity of the mutant library. Following PCR, each product was transformed into the yeast genome using a CRISPR-Cas9 gene-editing protocol targeting the stuffer sequence^42^. Once cells grew on selective media, cells from 4 petri dishes were pooled together to obtain 1 biological replicate, for a total of 4 biological replicates per fragment. We combined fragments 1-2-3, 3-4-5 and 5-6-7 to obtain the 3 final libraries. Unfortunately, after Illumina sequencing to validate our library, we noticed some mutants with low coverage. To obtain a more uniform library we combined our 4 biological replicates together. This resulted in a total of three libraries covering overlapping protein regions for each mating type.

### Screening for resistance to 5-FC

DMS libraries were screened in a bulk competition assay performed in duplicate (two separate culture tubes) for each of the three libraries (3 overlapping fragments each), for each mating type. We inoculated a 10 ml SC complete medium (NH_4_) preculture in a 50 ml tube using 150 ul of the glycerol stock for each library, for a total of 24 precultures. We grew them for 24 hours at 30°C with shaking in an orbital shaker at 200 rpm. The next day, we read the OD of each culture and washed 2 OD units with water to inoculate SC complete (NH_4_) + 0.00156 mg/ml 5-FC (estimated to inhibit approximately 50% of wild-type growth) and let them grow for 12 hours at 30°C at 200 rpm. We also kept 2 pellets of 5 OD units of those overnight cultures for DNA extraction (timepoint T0). After 12 hours, we repeated the same step, until we generated the cell pellets for timepoint T3. Cell pellets were stored at -80°C until DNA extraction. The number of mitotic generations for each sample was obtained by subtracting the logarithms of OD measurements after and before the 12-hour growth period between two timepoints and dividing by the logarithm of 2.

### DNA extraction and sequencing

Genomic DNA was extracted using a standard phenol/chloroform protocol, then quantified on NanoDrop (ThermoFisher) and run on agarose gels. Dilutions at 30 ng/µl were used for downstream PCR amplifications. The first PCR step added the Illumina PBS next to each fragment (see Supplementary Table 5 for primers). We had two versions of each PCR with different lengths of Ns (3 and 5 Ns) to generate diversity at the sequencing step. We combined primers so the length of Ns in the forward and reverse are always 8. We added 150 ng of DNA in each PCR reaction done with the Q5 polymerase (NEB). The PCR cycle was :

|  |  |  |
| --- | --- | --- |
| 1X | 98°C | 30sec |
| 5X | 94°C | 15sec |
|  | 50°C | 15sec |
|  | 72°C | 20sec |
| 15X | 98°C | 10sec |
|  | 65°C | 15sec |
|  | 72°C | 20sec |
| 1X | 72°C | 40sec |
| 1X | 22°C | 10sec |

PCR reaction from step 1 was checked on agarose gel and purified with magnetic beads (ratio 1:1, eluted in Tris 10 mM pH 8.0). To add Illumina indexes, we performed a second PCR step (see Supplementary Table 5 for primers). Purified PCR from step 1 were diluted at 0.1ng/µl. For the second step, we used 0.1ng of DNA with the Kapa polymerase (Roche) and this PCR cycle :

|  |  |  |
| --- | --- | --- |
| 1x | 98 °C | 3min |
| 18x | 98 °C | 30sec |
|  | 61 °C | 15sec |
|  | 72 °C | 35sec |
| 1x | 72 °C | 1min |
| 1x | 10 °C | Hold |

Each PCR reaction from step 2 was checked on agarose gel and purified with magnetic beads (ratio 1:1, elution in Tris 10 mM pH 8.0). Purified PCR from step 2 were quantified with the AccuClear Ultra High Sensitivity dsDNA Quantitation kit (Biotium). We generated the sequencing pool by combining 15 ng of each library. NGS was performed in PE300 on an Aviiti instrument at Plateforme d’analyse génomique (IBIS, Université Laval).

### DMS analysis

DMS sequencing data were analyzed using the gyōza DMS analysis pipeline v1.2.0^43^ to get selection coefficients for all variants for each combination of species, mating type and fragment. Briefly, variant read counts are converted into log2-fold change in allele frequencies between T0 and each post-screening time point. The values are then normalized with the median for silent mutants (excluding the wild-type nucleotide sequence because of its expected overabundance in the pools). A final normalization is applied by dividing by the number of mitotic generations to account for slightly different growth rates across conditions. Selection coefficients for each amino acid variant are obtained by first filtering out all alleles with fewer than 50 reads in any replicate, then calculating the median across synonymous codons. Finally, “fitness” estimates outputted by gyōza correspond to the median selection coefficient across replicates.

Plotting the selection coefficients at each time point showed that saturation was obtained at T2 (Supplementary Fig.14). Indeed, signal was steady at T1 and T2, then showed a sharp and global decrease at T3. This is likely caused by finite enrichment of advantageous mutants. Notably, this does not impact silent mutants, which confirms that the bias at T3 is not due to resistance-conferring secondary mutations outside the amplified locus. In downstream steps, we thus used only the selection coefficients calculated at T2.

The next step was to scale the data, by dividing with the median selection coefficient of nonsense mutants. Finally, we used linear regression on mutants in the overlaps between fragments F1/F2 and F2/F3 to normalize data across fragments, keeping the median value for each amino acid variant. We refer to the final score obtained as “resistance score” throughout, with values close to 0 indicating sensitivity and values close to 1 indicating 5-FC resistance.

In order to classify amino acid variants as either resistant or sensitive to 5-FC, we first calculated the average resistance score between the two mating types, then compared the obtained value to a threshold set 2 standard deviations below the mean resistance score of nonsense mutants. Variants with a resistance score above the threshold (ScCD, > 0.723; CnCD, > 0.553) were classified as resistant, while all others were classified as sensitive.

### Global epistasis model

To study epistasis, we took advantage of the replicability of our DMS experiment and considered the two mating types as individual replicates. The rationale was that comparable levels of experimental noise in both replicates would help us infer how much of the variance was driven by true specific mutational effects. In order to deconvolute specific from global epistasis, we fitted a global epistasis model to our DMS data using the multidms Python package^26^. The model assumes that mutations acted in additive fashion on an underlying, unobserved biological scale, termed the “latent phenotype”. This “latent phenotype” (*z*) is then mapped to the observed DMS data through a non-linear global transformation. In mathematical terms, we modelled the predicted resistance score (ŷ) with a sigmoid function (*g*):

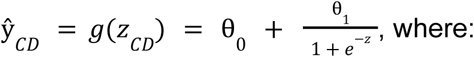

- ŷ is the predicted resistance score of any given mutant

- *z* quantifies the total latent phenotype of the mutant CD

- θ_0_ and θ_1_ are constant parameters, which we expect to approach 0 and 1, respectively, given how our resistance scores were normalized.

To model the impact of single amino acid substitutions, we define β as the mutational effect (compared to the corresponding wild-type) on this latent scale:

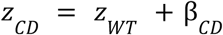

The expected resistance score of a mutant behaving similarly in both homologs is dictated purely by the global sigmoid function, where the mutation contributes an identical β to the latent phenotype (*z*) of both homologs. On the other hand, epistatic mutants are characterized by a non-null difference in predicted mutational effects, which can be expressed as a function of a “shift” (Δ) in the latent space:

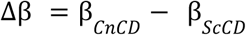

This shift (Δβ) explicitly quantifies the magnitude of specific epistasis. The next step is to estimate the model parameters, which multidms achieves using an objective function combining two terms to balance accuracy with biological interpretability:

- A loss term that corresponds to the difference between predicted (ŷ) and experimentally derived (y) resistance scores to maximize accuracy

- A LASSO regularization term that shrinks marginal shifts to 0 by applying a penalty proportional to |Δβ|

In other words, if an inferred epistatic shift is weakly supported by the experimental data, Δ is forced to 0. This approach ensures a majority of true epistatic mutations are identified, consistent with our expectations and previous studies.

### Classification of epistatic mutations

Mutations were classified as epistatic based on their epistatic shift (Δβ). Since nonsense mutants have a resistance score of 1 on average regardless of the species, the distribution of their shift should be a normal distribution centered around 0. We therefore generated a null distribution from 10,000 bootstrap replicates. For each missense mutant, we calculated an empirical p-value corresponding to the proportion of bootstrap replicates with |Δβ_nonsense_| > |Δβ_missense_|. Mutations were classified as epistatic if the null hypothesis could be rejected at a 0.1% false discovery rate (using Benjamini-Hochberg correction) or non-epistatic if the null hypothesis could not be rejected. Finally, since the distribution of the differences in resistance scores was strongly bimodal, we re-classified as non-epistatic the few significant mutants for which this value was between -0.2 and 0.2.

### Features

We used the 3D structures to estimate the effects of amino acid substitutions on protein stability and interaction using FoldX^44^. The RepairPDB function was run 10 times to repair incorrect torsion angles, Van der Waals clashes and total energy of the structure. We used the MutateX Python package to perform an in silico DMS of the entire protein. Since the protein forms a homodimer, mutations were introduced symmetrically in both chains. The BuildModel function from FoldX was applied to compute the change in Gibbs free energy (ΔΔG) between each mutant and the wild type, while the AnalyseComplex function was used to evaluate the effect of mutations on the interaction energy between the two subunits.

We used GEMME^45^ and ESM1b^46^ to predict the functional impact of mutations. To run GEMME, we first produced a multiple sequence alignment of CD orthologous sequences. This was obtained through iterative sequence search by using the protein sequence of ScCD as the query for the HHblits function from the HH-suite3^47^. Initial filtering criteria included a minimum coverage of 80% of the length of the query sequence, between 20% and 98% identity with the query, while the e-value cutoff for inclusion was 0.0001. These filtering criteria were then gradually relaxed to get a minimum of 200 sequences in the final alignment. GEMME was then run locally to predict the effect of every single amino acid substitution in the query. The same approach (HHblits, followed by GEMME) was repeated, this time using the protein sequence of CnCD as the query. We computed ESM scores using the ESM1b-derived protein language model variant described by Brandes et al^24^. For each missense mutation, the variant effect score was computed as the difference in log-likelihood between the mutant and the wild-type amino acid at the same position, for both proteins.

We computed the distance between each amino acid residue and the zinc atom, which we used as a reference point because it marks the active site. We also generated the contact matrices with custom Python scripts using BioPython’s PDBParser. Residues were considered in contact with one another if they were less than 5 Å away. From these matrices, we calculated the proportion of conserved contacts for each residue, defined as the ratio between the number of shared contacts (present in both contact matrices) and the total number of unique contacts for that residue.

Residue flexibility was assessed with a custom Python script by comparing the positions of Cα atoms between the open and closed states of the subunit, after structural alignment with Chimera’s MatchMaker tool. Relative solvent accessibility (RSA) was computed using freeSASA, while residue buriedness was obtained with a script available at https://github.com/rodogi/buriedness. Buriedness is conceptually similar to RSA but does not reach null values, allowing differentiation of residues that remain shielded from solvent. Secondary structure and backbone torsion angles were predicted with STRIDE (http://webclu.bio.wzw.tum.de/stride/).

Amino acid physicochemical properties were obtained from ProtScale (ExPASy; https://web.expasy.org/protscale/). To reduce redundancy among correlated features, we performed hierarchical clustering of amino acid properties and selected the most representative feature from each cluster. For model training, each property was associated with both the wild-type and the mutant residue, and the difference (Δ property) was computed.

### Machine learning

Machine learning models were implemented using a standardized pipeline composed of three steps: (i) feature scaling with *StandardScaler*, (ii) optional class balancing through resampling (SMOTE, random oversampling, random undersampling, ADASYN or no resampling) and (iii) model fitting with one of several classifiers (logistic regression, decision tree, support vector machine, random forest, balanced random forest, XGBoost, or CatBoost). For each classifier, hyperparameters were optimized through randomized search over predefined parameter distributions, including both model-specific variables and the resampling strategy. RandomizedSearchCV was performed using 200 iterations and a repeated stratified 5-fold cross-validation scheme (5 folds × 3 repeats). Multiple evaluation metrics were computed during tuning, including the Matthews correlation coefficient (MCC), balanced accuracy, precision, and recall. MCC was designated as the primary metric for model selection (refit criterion). The best-performing models were retrained on the full training set and assessed on the test set. The best model was then used to predict the ortholog dataset.

### Western blot

A N-terminal 1XFLAG was fused to *FCY1* by PCR by amplifying the gene from the genomic DNA of strains AKD1139 and AKD1233 for ScCD and CnCD, respectively, using primers CLOP351-A3, CLOP351-C3 and CLOP216-G11. The resulting PCR products were integrated in strains AKD1015 and AKD1021 for ScCD and CnCD, respectively, using CRISPR-Cas9 to target the stuffer sequence at the *FCY1* locus. The Cas9-encoding plasmid was lost by culturing cells from individual colonies overnight in YPD medium. Strains were then spotted on the appropriate antibiotics to confirm loss of the marker. Correct genomic integration and FLAG labeling were confirmed by colony PCR using primers CLOP326-A11/CLOP314-E8. The resulting strains, AKD1463, AKD1464, AKD1466 and AKD1467, were used for the Western blot.

The four strains and a negative control expressing an unflagged Sc*FCY1* (AKD1139) were grown overnight in SC (MSG) (0.174% yeast nitrogen base without amino acids, 2% glucose, standard drop-out mix, 0.1% monosodium glutamate) in triplicates. The next day, strains were diluted to 0.15 OD/ml in 15 mL fresh SC (MSG) and grown until they reached ∼ 1 OD/ml. The cultures were spun down at 21,100 x *g* for 5 min. Supernatant was discarded and cell pellets washed in 1 mL sterile water and transferred in 1.5 mL tubes. Cells were spun down for 1min30 at 1,000 x *g*. Water was removed by aspiration and cell pellets were frozen at -80°C.

Cell pellets were thawed at room temperature and resuspended in Complete Mini protease inhibitor mix (Roche 11836153001) at 0.05 OD/ul concentration. Cell suspension (200 μL) was mixed with glass beads and put on a glass beater for 5 min. SDS 10% was added to a final concentration of 1% (20 μL in 200 μL) and cell suspension was boiled for 10 min. Suspension was cleared of cell debris by centrifugation for 5 min at 15,000 rpm. Cell lysates were separated by SDS-PAGE on a 15% acrylamide gel (10 uL cell lysate, 7.5 μL water, 2.5 μL DTT 1M and 5 μL LB5X) for 50 min at 175 V. Proteins were transferred to a nitrocellulose membrane with a Semi-dry system (Amersham TE 77 PWR). Proper protein loading and transfer was confirmed with Ponceau Red staining of the membrane. Membrane was blocked overnight in blocking buffer from LI-COR (#927-70003). The next day, the membrane was incubated for 1 h with Anti-FLAG M2 antibody (0.6 µg/ml final, Sigma F3165) at room temperature. Membrane was washed 3 times 5 min with PBS-T (Na_2_HPO_4_ 81 mM, NaH_2_PO_4_ 24.7 mM, NaCl 100 mM). Membrane was incubated for 1h with the Anti-mouse 800 antibody (0.075 µg/ml final) at room temperature. Imaging of the membrane was done with a LI-COR system (Odyssey XF) with a 30 seconds scan in channel 700 and a 2 min scan in channel 800.

### Protein thermal stability assays

Proteins were expressed and purified according to the protocol described in the *E. coli* expression and protein purification section. Protein thermal stability was assessed using nano-differential scanning fluorometry (nanoDSF) on a Prometheus NT.48 instrument (NanoTemper Technologies). Purified recombinant protein samples (10 µL), were loaded into nanoDSF Grade Standard Capillaries. Thermal unfolding was monitored over a temperature range of 20 °C to 95 °C with a 1 °C/min ramp. Melting temperatures (T_m_) were determined from the maximum of the first derivative of the 350/330 nm fluorescence ratio using PR.ThermControl software (version 2.3.1). T_m_ values were reported only for samples showing a dominant transition in the first derivative curve. Samples without a clearly defined peak were considered not to have a measurable T_m_ under the experimental conditions. All protein variants were analyzed in triplicate.

### In vitro protein activity assays

CnCD and ScCD wild-type and mutant activities were assessed by monitoring the conversion of substrate 5-FC into 5-FU, measured as a shift in absorbance at 237 nm using a Corning UV transparent 96-well (#3635). For each assay, 100 μl of purified recombinant protein at a concentration of 0.01 mg/ml was mixed with equal volume of 5-FC (1 mM) in triplicates. Absorbance was measured every 20 seconds over 30 minutes for ScCD proteins and over 75 minutes for CnCD proteins. All measurements were performed in triplicates using a Biotek Synergy H1 (Agilent technologies) plate reader with the temperature set at 25°C.

## Acknowledgments

Part of the research described in this paper was performed using beamlines CMCF-BM and CMCF-ID at the Canadian Light Source, a national research facility of the University of Saskatchewan, which is supported by the Canada Foundation for Innovation (CFI), the Natural Sciences and Engineering Research Council (NSERC), the National Research Council (NRC), the Canadian Institutes of Health Research (CIHR), the Government of Saskatchewan, and the University of Saskatchewan.This research used resources of the Advanced Photon Source; a U.S. Department of Energy (DOE) Office of Science User Facility operated for the DOE Office of Science by Argonne National Laboratory under Contract No. DE-AC02-06CH11357. Use of the Lilly Research Laboratories Collaborative Access Team (LRL-CAT) beamline at Sector 31 of the Advanced Photon Source was provided by Eli Lilly and Company, which operates the facility.

We thank members of the Landry and Shi labs for critical feedback.

## Authors contributions

Obtained funding: CRL and RS; Designed research: CRL, RS, MEP, AKD; Performed experiments: AKD, MEP, PCD, EA, JG; Data analysis and interpretation: RD, AP, MEP, SD, RS, CRL; Wrote the initial draft of the manuscript: CRL, RD, AP, AKD, MEP; Editing and polishing: MEP; RD; AKD; SD, AP, PCD, RS and CRL.

## Extended data figures

**Extended Data Figure 1.**
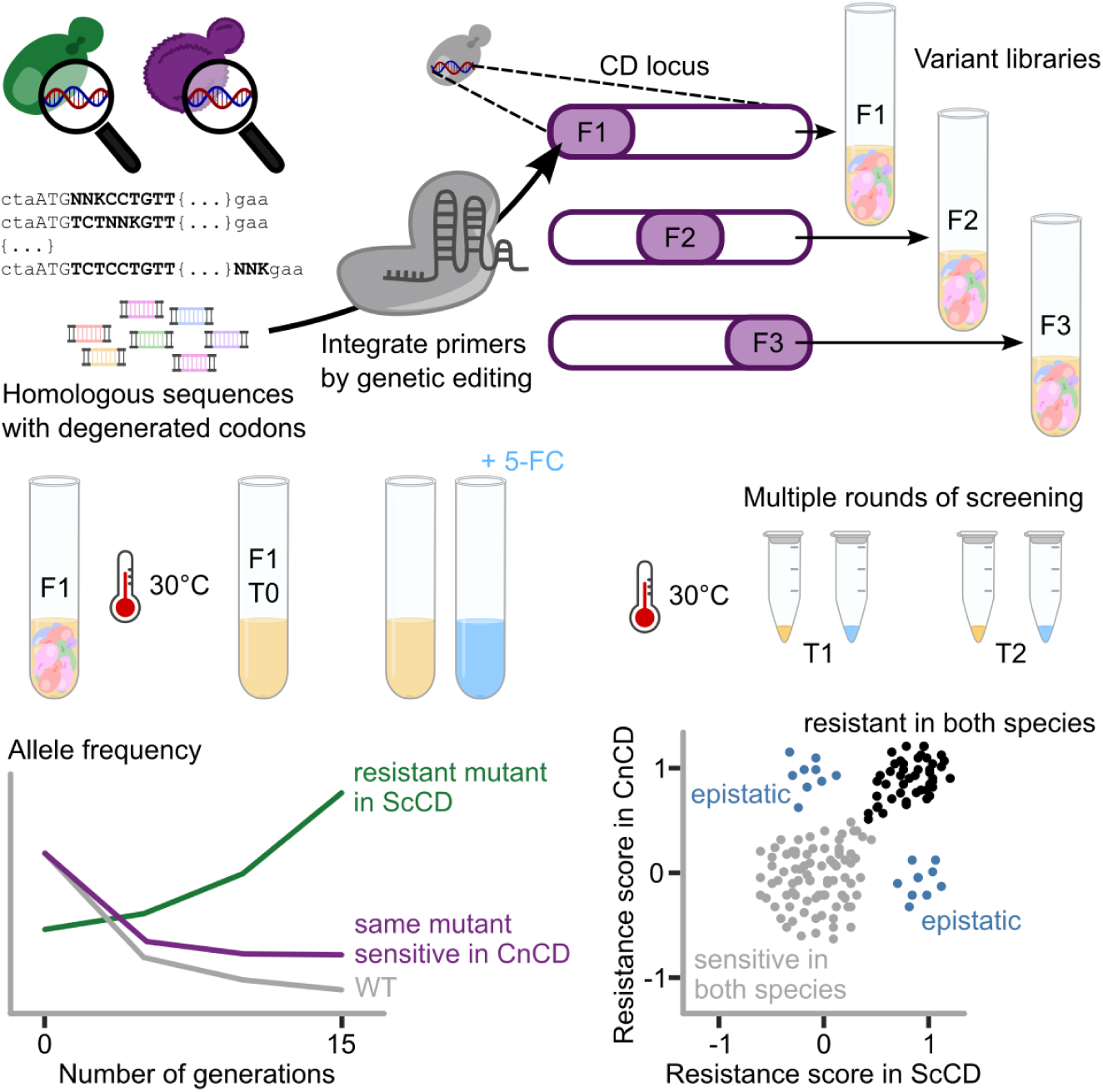
Experimental design. Schematic representation of the deep-mutational scanning experimental and computational design. Libraries of ScCD and CnCD (with degenerate codons at each position of overlapping fragments) were integrated at the genomic locus of *FCY1* in haploid strains of *S. cerevisiae*. For each fragment separately, libraries were expressed by culturing overnight in rich medium. Cultures were passaged every 5 generations for a total of 15 generations in medium with or without 5-FC. At each time point, cells were pelleted and amplicon sequencing was performed from genomic DNA to measure the relative allele frequency of each variant. Resistance scores can be obtained from relative changes in allele frequencies. From there, one can identify mutants resistant or sensitive to 5-FC in both backgrounds, as well as mutants showing background-dependent effects, here labeled as “epistatic”.

**Extended Data Figure 2.**
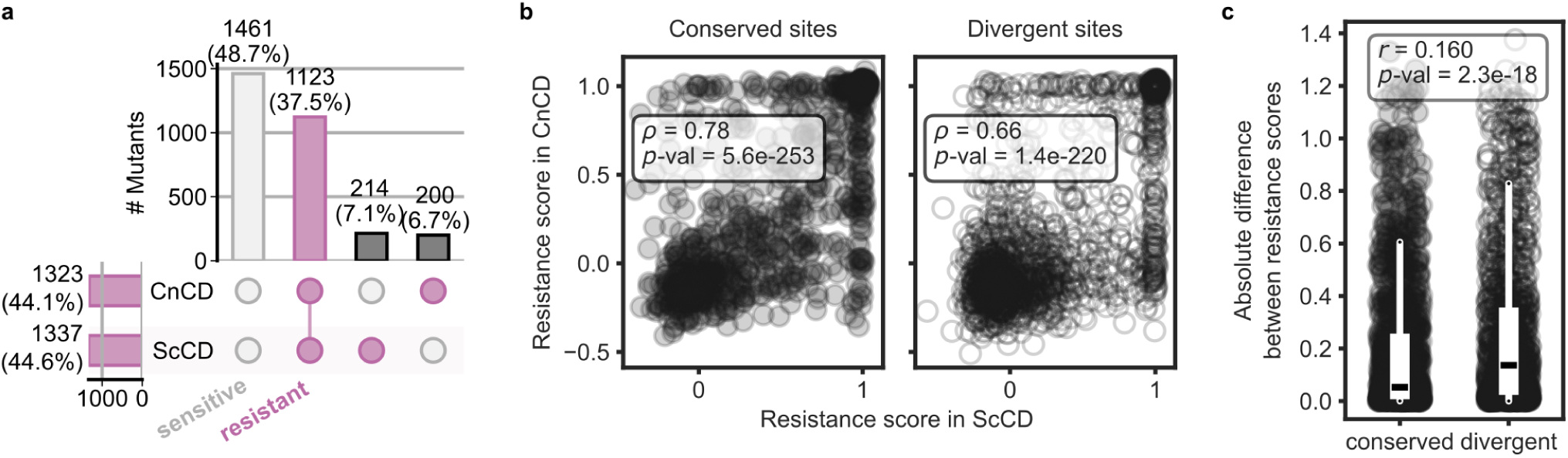
Correlation between fitness effects of missense mutants in ScCD and CnCD at conserved and divergent sites. (**a**) UpsetPlot indicating the number (and proportion) of single amino acid substitutions classified as resistant (pink) or sensitive (gray) in ScCD, CnCD or both. To prevent any redundancy in the dataset, we excluded substitutions introducing the wild-type residue from either species. Additionally, we excluded any substitution for which we could not have the classified equivalent in the other species, to make sure missing data would not be miscounted as sensitive mutants. (**b**) Scatter plot of resistance scores for mutants at conserved (left) and divergent sites (right). For each panel, the corresponding Spearman correlation coefficient and *p*-value are indicated. (**c**) The absolute differences between resistance scores for mutants at conserved sites and divergent sites were compared using a Mann-Whitney (two-sided) test. The effect size (*r*) and *p*-value are indicated. The same dataset was used for all three panels.

**Extended Data Figure 3.**
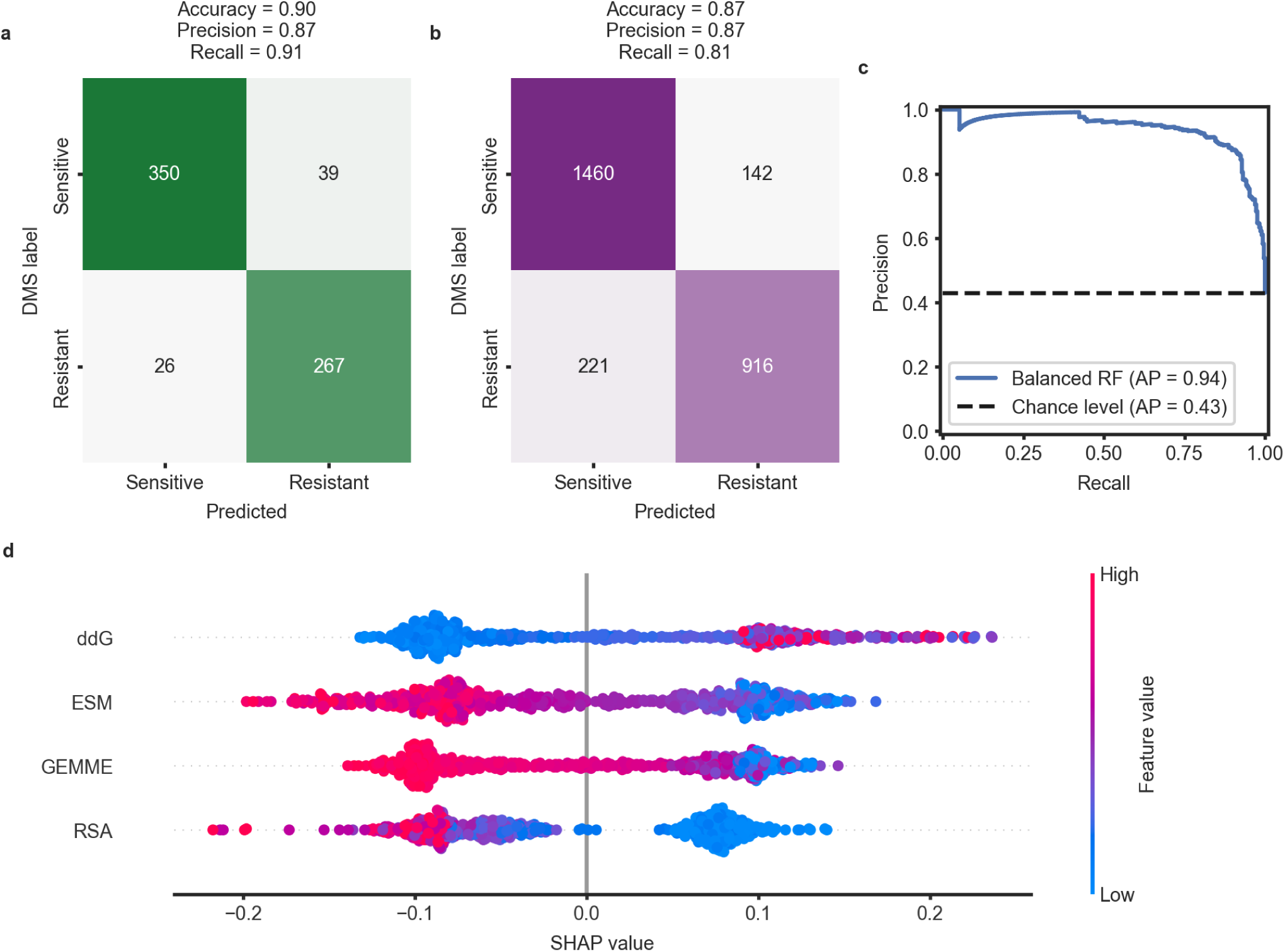
Predictions of 5-FC resistance in CnCD. A balanced random forest model was trained using single amino acid substitutions classified as resistant or sensitive to 5-FC in ScCD. The model was used to predict resistance for the equivalent mutants in CnCD. Classified mutants of CnCD from our DMS experiment were used to validate the predictions. Panels (**a**) and (**b**) show the metrics and confusion matrices summarizing prediction performance. Panel (**a**) reports performance on the held-out test set from the ScCD training data, whereas panel (**b**) presents model performance when predicting resistance in the orthologous CnCD dataset. Panel (**c**) displays the ROC curve computed on the held-out test set. SHAP values represented in panel (**d**) indicate which model features impacted the most the decision of classifying any mutant. Only the first four are displayed, by order of contribution. For example, a high ΔΔG predicted by FoldX indicates structural destabilization and shifts the model toward predicting resistance.

## Data and materials availability

All data are available in the main text or the supplementary materials. Sequencing data are available at the NCBI Sequence Read Archive (SRA) under BioProject PRJNA1333547: http://www.ncbi.nlm.nih.gov/bioproject/1333547. The coordinates and structure factors of the have been deposited on the RCSB Protein Data Bank (CnCD : 36FA, AnCD: 36FB, NgCD: 36FC, CnCD_A99S: 36FD, ScCD_M100T: 36GM, CnCD_G120E: 36FE, CnCD_G119_G120insK: 36FF, CnCD_G119_G120insK bound to 4-oxidanyl-3,4-dihydro-1H-pyrimidin-2-one: 36FG).

## Supplementary material

**Supplementary Figure 1.**
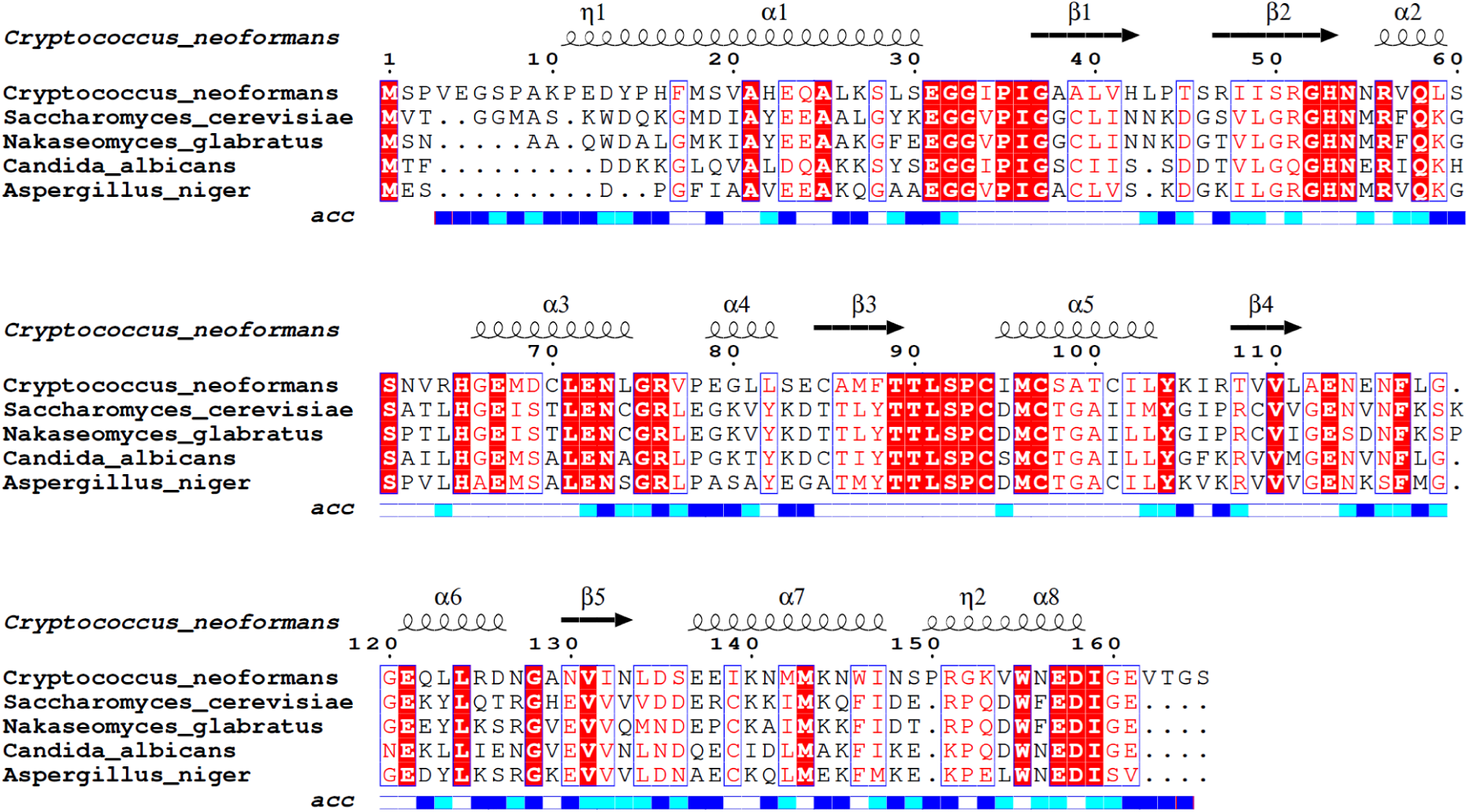
Multiple sequence alignment of 4 CD homologs. Protein sequences were aligned using MUSCLE v5^1^. The alignment was visualized with ESPript 3.0^2^, applying the BLOSUM62 similarity coloring scheme. Secondary-structure elements displayed above the alignment correspond to those from the *C. neoformans* CnCD crystal structure (PDB 36FA), with α-helices shown as cylinders and β-strands as arrows, as generated automatically by ESPript. Conserved residues are highlighted according to ESPript defaults, with strictly conserved positions shown in red boxes and similar residues in red text.

**Supplementary Figure 2.**
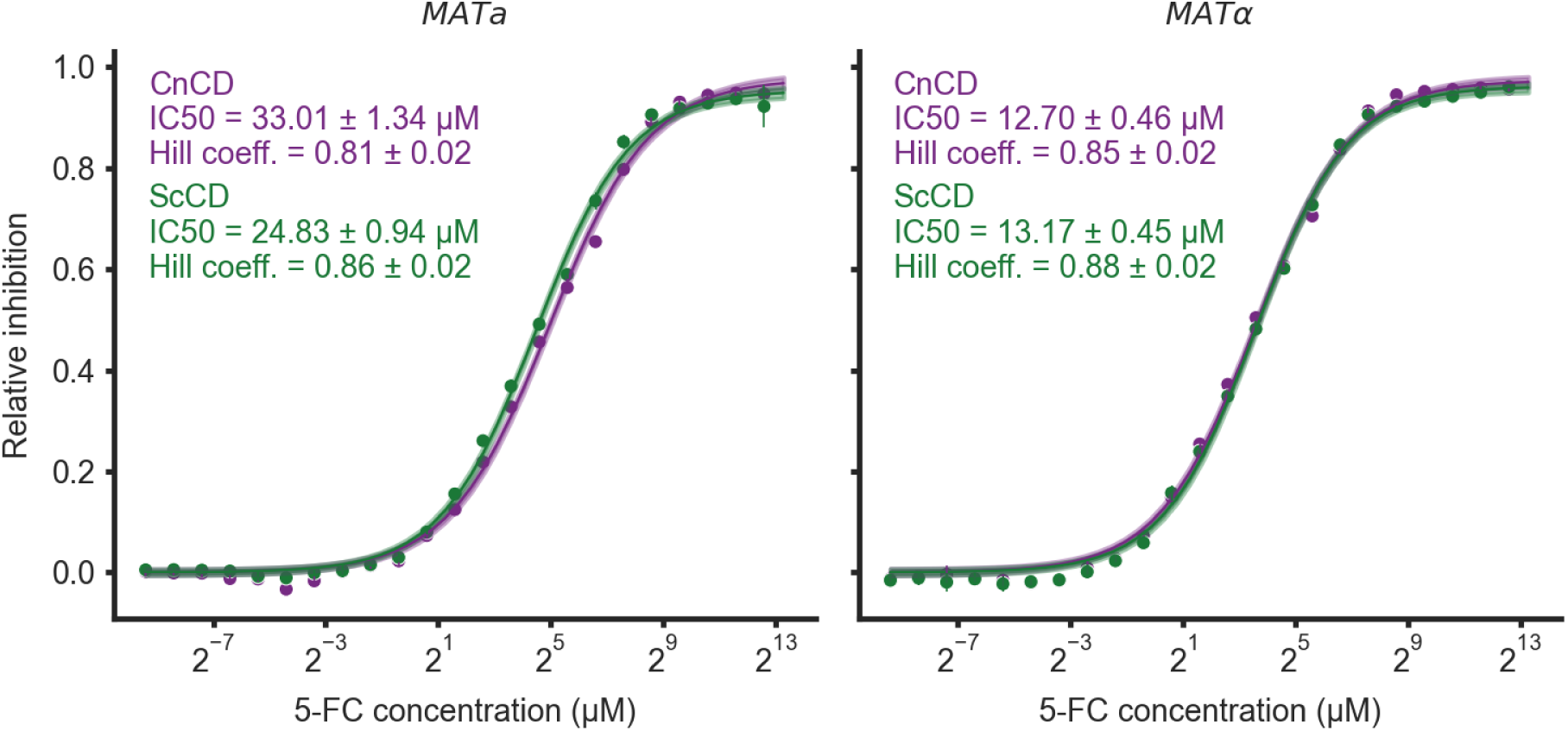
Dose-inhibition curves of *Saccharomyces cerevisiae* expressing ScCD and CnCD in a gradient of concentration of 5-FC. Inhibition curves of *S. cerevisiae MATa* (left) or *MATɑ* (right) expressing ScCD (green) or CnCD (purple) were obtained by growing the strains for 24h at 30°C using 24 different concentrations of 5-FC (0 μg/mL and twenty-three 2-fold dilutions starting at 800 μg/mL). The condition without 5-FC was used to normalize maximum growth rates obtained from four biological replicates. The log logistic function was used to model inhibition coefficients (1 - growth rate). Curve fitting and confidence bands were determined via the Delta Method using the model covariance matrix. The IC50 (concentration at which 50% of growth is inhibited) was obtained by plugging the parameters estimated by the model back into the log logistic equation. The slope indicated on each panel, here referred to as a Hill coefficient for simplicity, corresponds to one of the estimated model parameters. Data points represent the mean across replicates, while error bars denote the 95% confidence interval calculated as ± 1.96 (t-statistic) multiplied by the standard error of the mean.

**Supplementary Figure 3.**
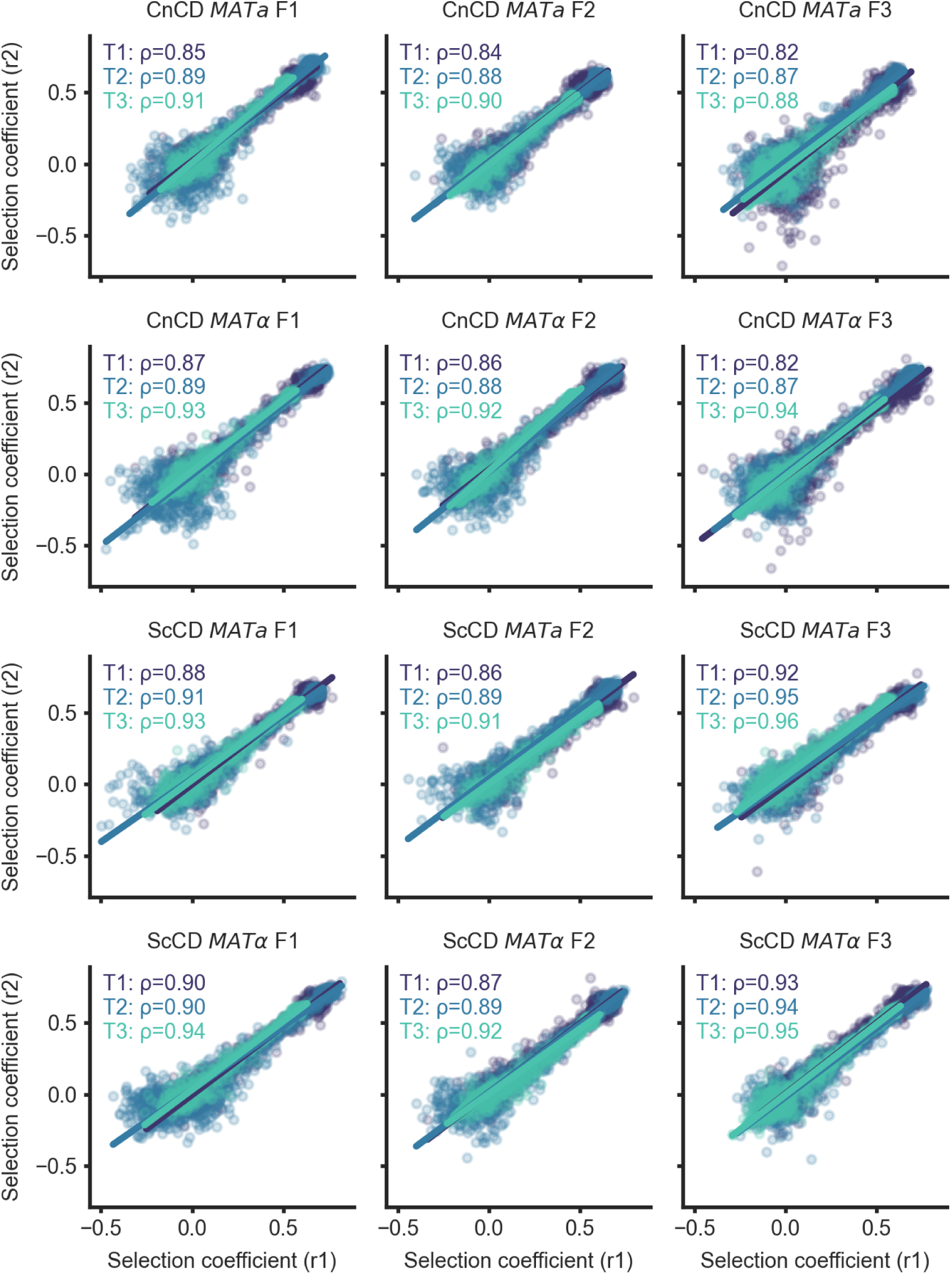
Replicability of the DMS experiment through time. Scatter plots showing the selection coefficients of replicates 1 (r1) and 2 (r2) (independent growth cultures and all downstream steps) for each construction. The two-mating type strains represent libraries independently constructed. Time points at which cells were sampled are indicated by the different shades of blue. The two mating types represent two independent strain library constructions. The Spearman correlation coefficient is reported for each time point and condition.

**Supplementary Figure 4.**
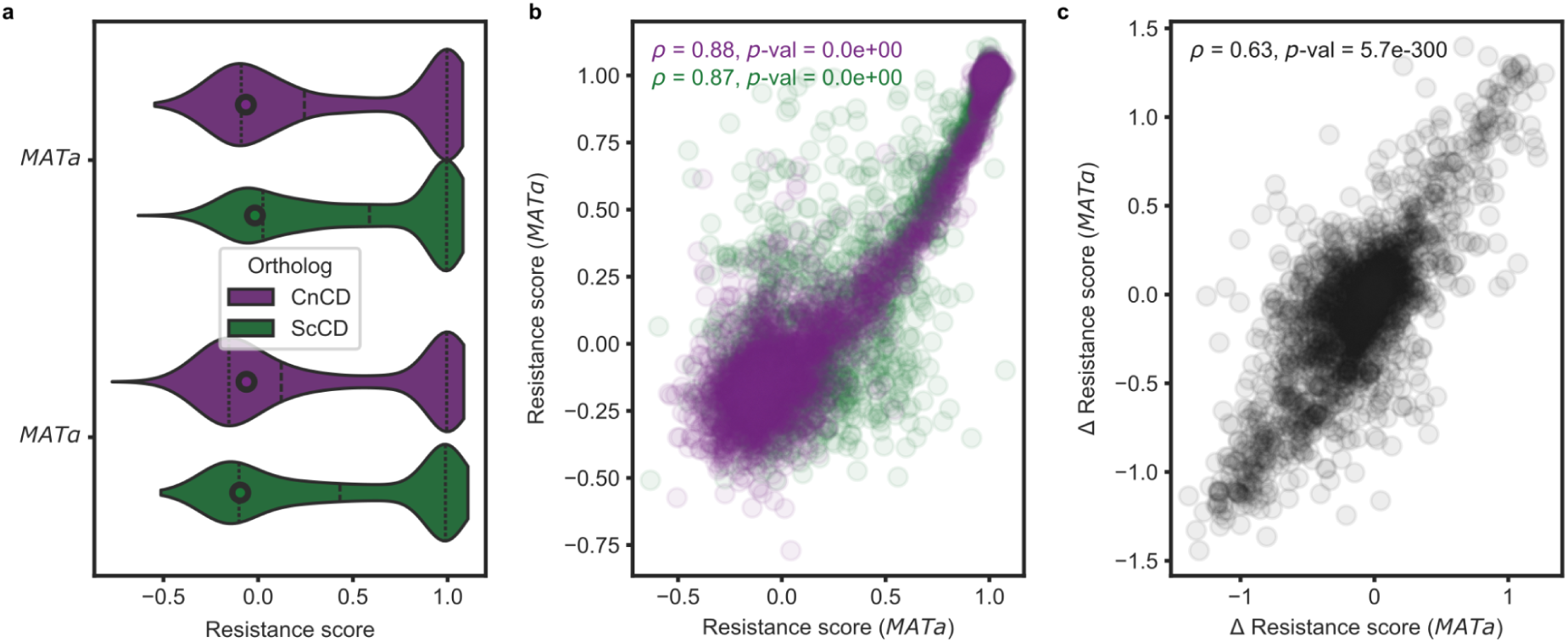
Replicability of the DMS experiment using two haploid backgrounds for library construction. (**a**) Distribution of resistance scores for each ortholog across the *S. cerevisiae* mating types. Wild-type scores are indicated by black circles. (**b**) Correlation of resistance scores between mating types for CnCD (purple) and ScCD (green), with Spearman correlation coefficient and *p*-value indicated. (**c**) Correlation between the differences in scores of matching mating types (CnCD - ScCD), with Spearman correlation coefficient and *p*-value indicated. In this last panel, nucleotide substitutions that introduced the wild-type residue from either species were excluded. In all three panels, resistance scores correspond to selection coefficients scaled between 0 (silent) and 1 (nonsense).

**Supplementary Figure 5.**
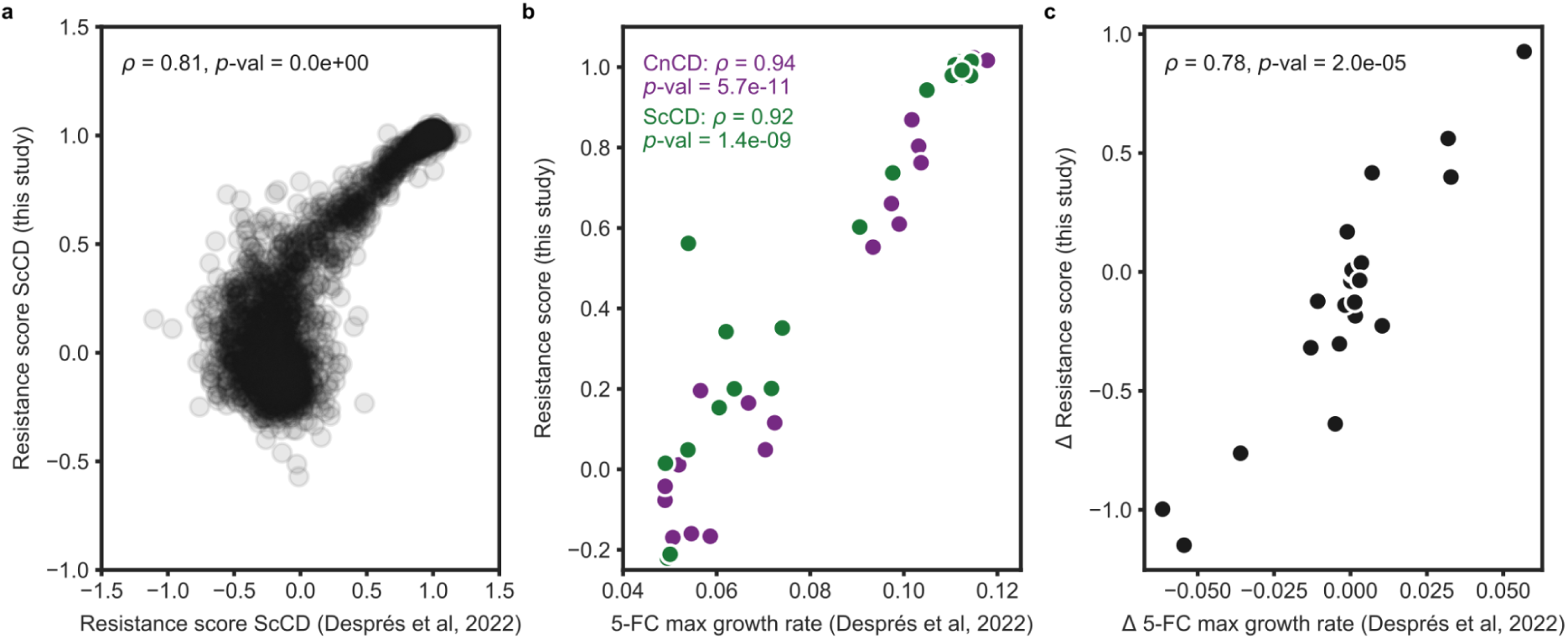
Correlation between the data obtained in this paper and those from a previous experiment. (**a**) Scatter plot showing the correlation between resistance scores of ScCD obtained here and the DMS dataset of Després et al (2022)^3^. For both, we used the same libraries, but they were screened independently, then pooled differently for sequencing and finally, the scores were normalized differently. (**b**) Scatter plot showing the correlation between the maximum growth rate in media containing 5-FC (mean of two biological replicates) for 22 individually re-constructed mutants from Després et al (2022)^3^ and their resistance score estimated in this study. (**c**) Scatter plot showing the correlation between the differences in resistance score obtained in this study and in maximum growth rate in media containing 5-FC for the 22 matched mutants examined in both studies (derived from the data in panel b). In all panels, the corresponding Spearman correlation coefficients and *p*-values are indicated.

**Supplementary Figure 6.**
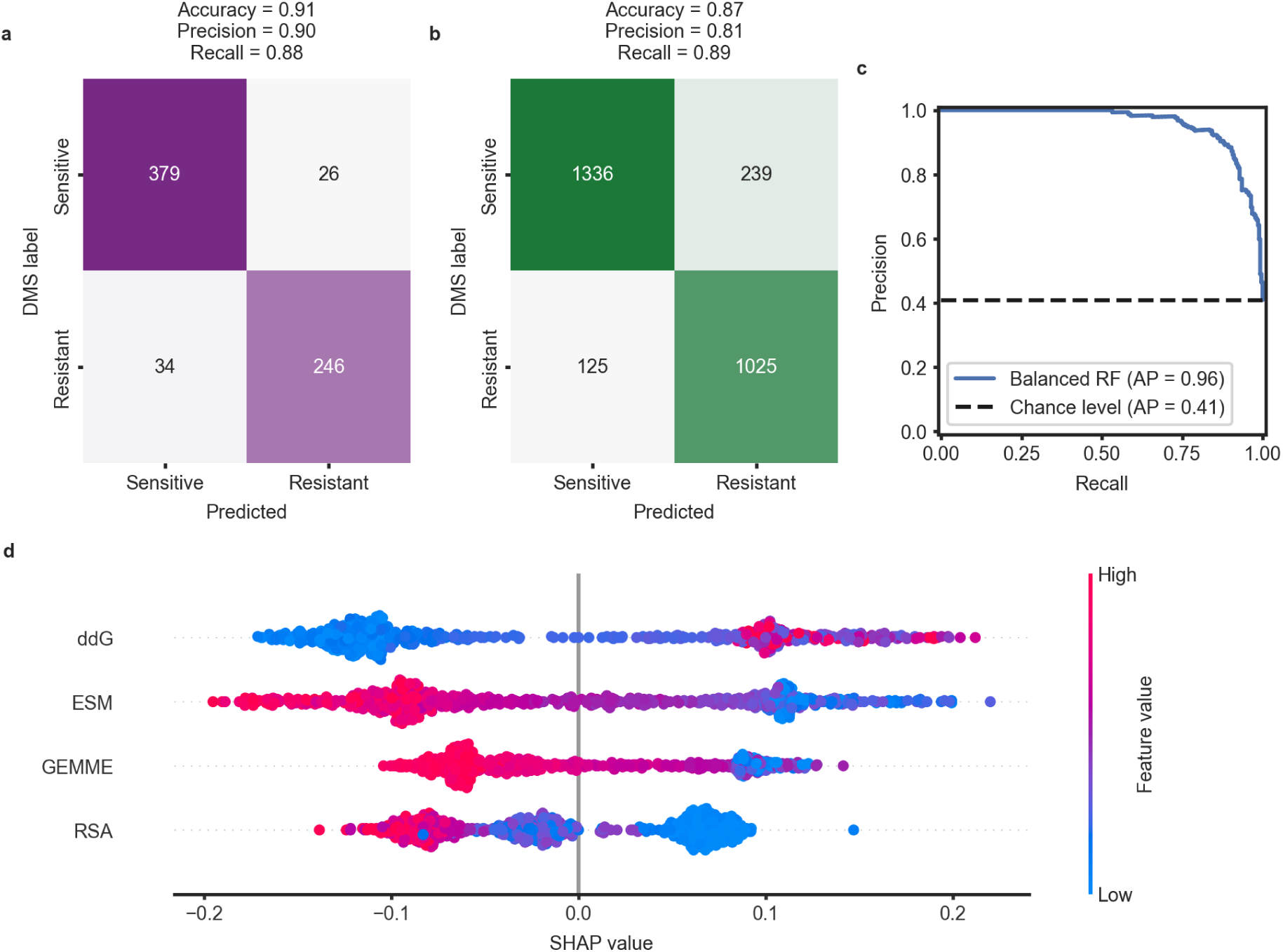
Predictions of resistance to 5-FC caused by mutations in ScCD. A balanced random forest model was trained on single-amino-acid substitutions classified as resistant or sensitive to 5-FC in CnCD. The model was used to predict resistance for the equivalent mutants in ScCD. Experimentally classified ScCD mutants from our DMS experiment were used to validate the predictions. Panels (**a**) and (**b**) show the metrics and confusion matrices summarizing prediction performance. Panel (**a**) reports performance on the held-out test set from the CnCD training data and (b) presents model performance when predicting resistance in the orthologous ScCD dataset. Panel (**c**) displays the ROC curve computed on the held-out test set. SHAP values represented in panel (**d**) indicate which model features had the most impact on the decision of classifying any mutant. Only the first four are displayed, in order of contribution. For example, a high ΔΔG predicted by FoldX indicates structural destabilization and shifts the model toward predicting resistance.

**Supplementary Figure 7.**
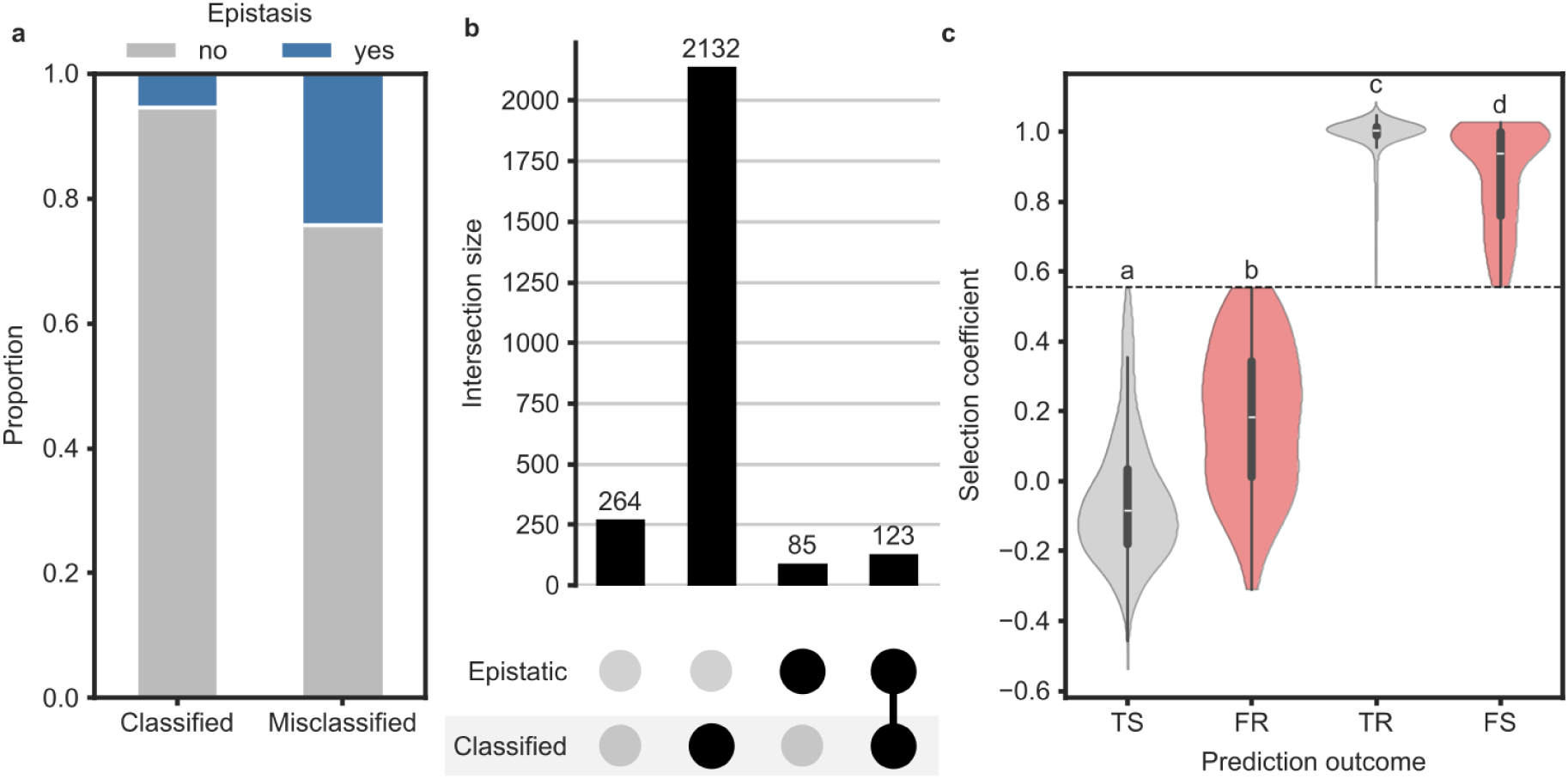
Relationship between epistasis and model misclassification. (**a**) Proportion of epistatic versus non-epistatic mutations among correctly classified and misclassified variants. Epistatic mutations are those that lead to different phenotypes in the two orthologs (resistance or susceptibility). Misclassified variants show an enrichment for epistasis compared to correctly classified ones. Epistatic mutations were significantly enriched among misclassified variants (24.3%, 85/350) compared with correctly classified variants (5.5%, 123/2254; χ² = 143.58, p < 0.001) (**b**) UpSet plot showing the size of intersections between epistatic mutations and classified/misclassified variants. (c) Distribution of experimentally measured selection coefficients for mutations grouped by prediction outcome: true sensitive (TS), false resistant (FR), true resistant (TR), and false sensitive (FS). Violin plots show the distribution of selection coefficients, with embedded boxplots indicating the median and interquartile range. Misclassified variants (FR and FS) are highlighted in red and tend to exhibit less extreme selection coefficients than correctly classified variants. Differences among groups were significant (Kruskal–Wallis test: H = 2066.78, p < 0.001). Post hoc Dunn tests indicated significant differences among all pairwise comparisons (p < 0.001); different letters above violins denote statistically distinct groups.

**Supplementary Figure 8.**
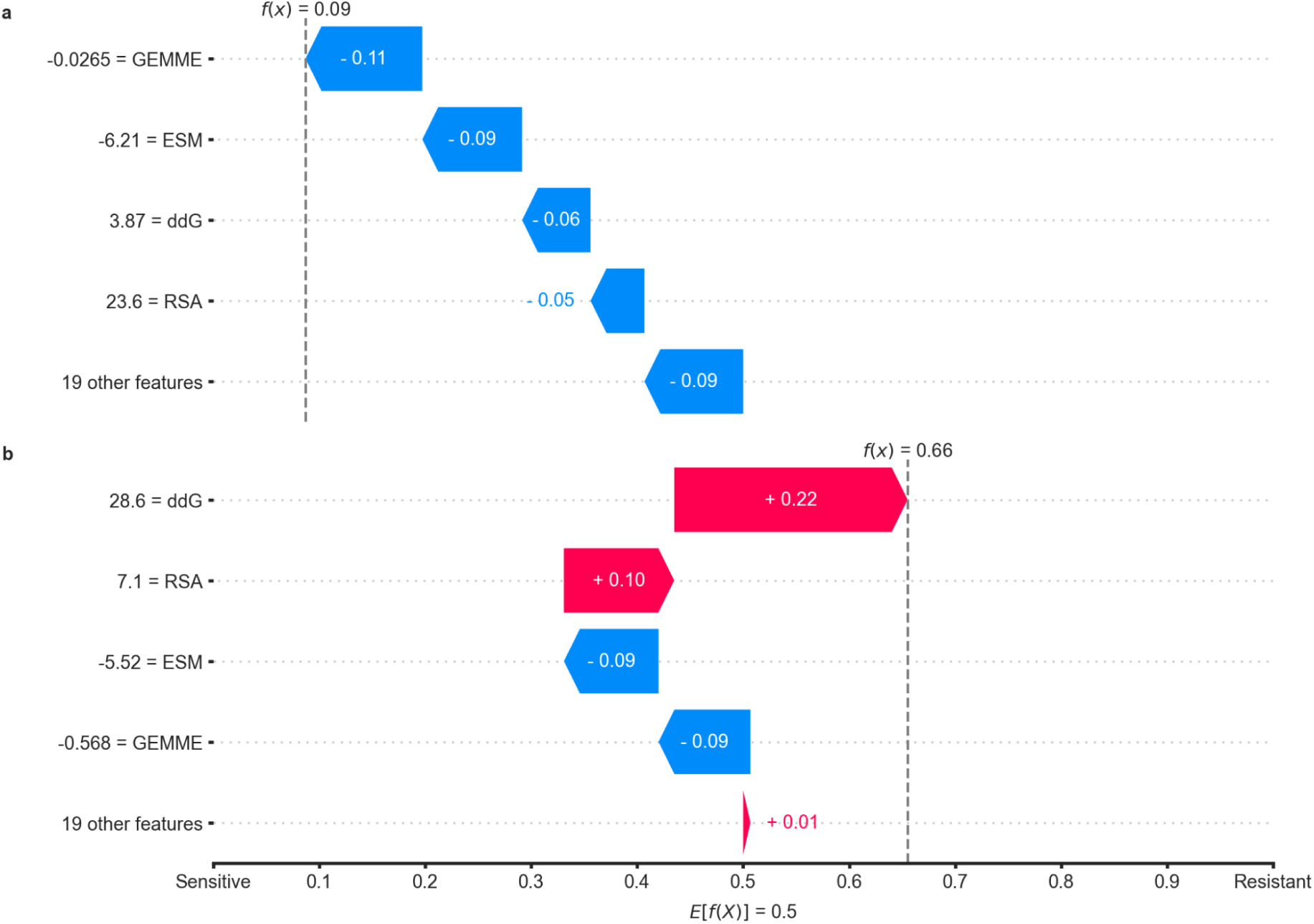
SHAP explanations for variants in which substitution to the homologous wild-type residue results in resistance (loss of function). SHAP waterfall plots illustrating feature contributions to model predictions for two representative variants from the subset in which substitution to the homologous wild-type residue leads to loss of function. Red bars indicate features that increase the predicted probability of resistance, whereas blue bars indicate features that increase the predicted probability of sensitivity. Feature values are shown on the left of each plot. (**a**) Example of a misclassified variant (L43N). Evolutionary features (GEMME and ESM) favor the homologous wild-type residue and drive the prediction toward sensitivity. As a result, the variant is incorrectly predicted as sensitive. (**b**) Example of a correctly classified variant (G152P). The large positive contribution of the ΔΔG feature, together with structural context captured by relative solvent accessibility (RSA), outweighs the opposing influence of evolutionary features and drives the prediction toward resistance.

**Supplementary Figure 9.**
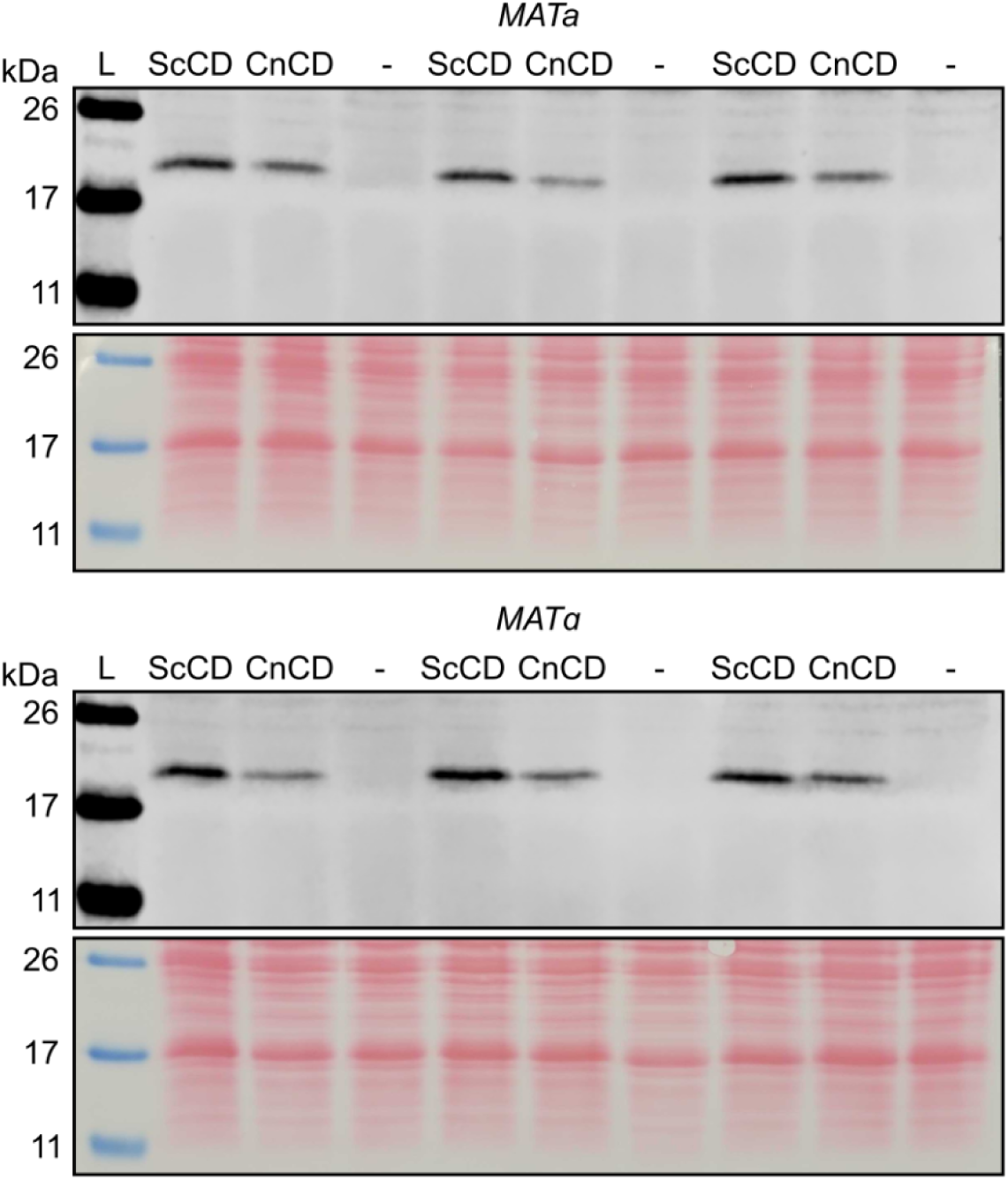
Abundance of CDs expressed in *S. cerevisiae*. Yeast cells were grown until the exponential phase in SC (MSG). Western blotting was performed using an antibody directed against the FLAG (Sigma product F3165) tag fused to the N-terminal of CDs. Tracks are annotated with what was loaded (in three replicates, each in the same order), including the ladder (L) and a negative control (-). The negative control was the parental strain without the FLAG tag (AKD1139). Panels at the bottom for each mating type show the result of Red Ponceau staining, used as loading and transfer control. The four strains used with the FLAG tag are AKD1463 (his3Δ1 leu2Δ0 met15Δ0 ura3Δ0 FLAG-FCY1opt Scer ho::NATNT2 MATa), AKD1464 (his3Δ1 leu2Δ0 met15Δ0 ura3Δ0 FLAG-FCY1opt Cneo ho::NATNT2 MATa), AKD1466 (his3Δ1 leu2Δ0 lys2Δ0 ura3Δ0 FLAG-FCY1opt Scer ho::HPHNT1 MATalpha) and AKD1467 (his3Δ1 leu2Δ0 lys2Δ0 ura3Δ0 FLAG-FCY1opt Cneo ho::HPHNT1 MATalpha).

**Supplementary Figure 10.**
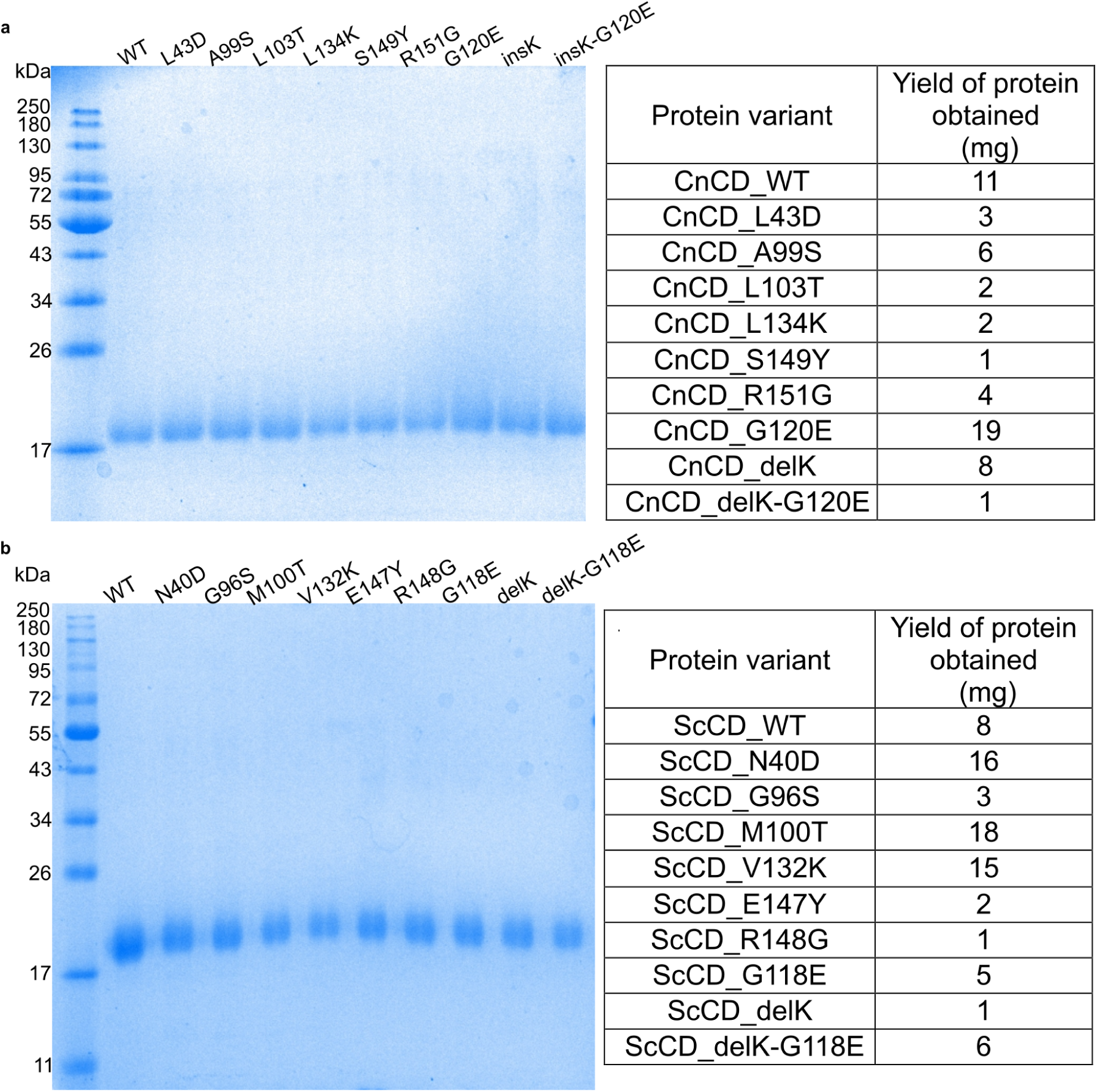
SDS-PAGE analysis and final yields of purified CD proteins. (**a**) CnCD and (**b**) ScCD WT or variant proteins after cobalt affinity chromatography and buffer exchange into the final buffer. Proteins were resolved on a 12.5% acrylamide gel (200V, 45 min) and stained with Coomassie Blue R-250. The accompanying tables report the final protein yield for each preparation.

**Supplementary Figure 11.**
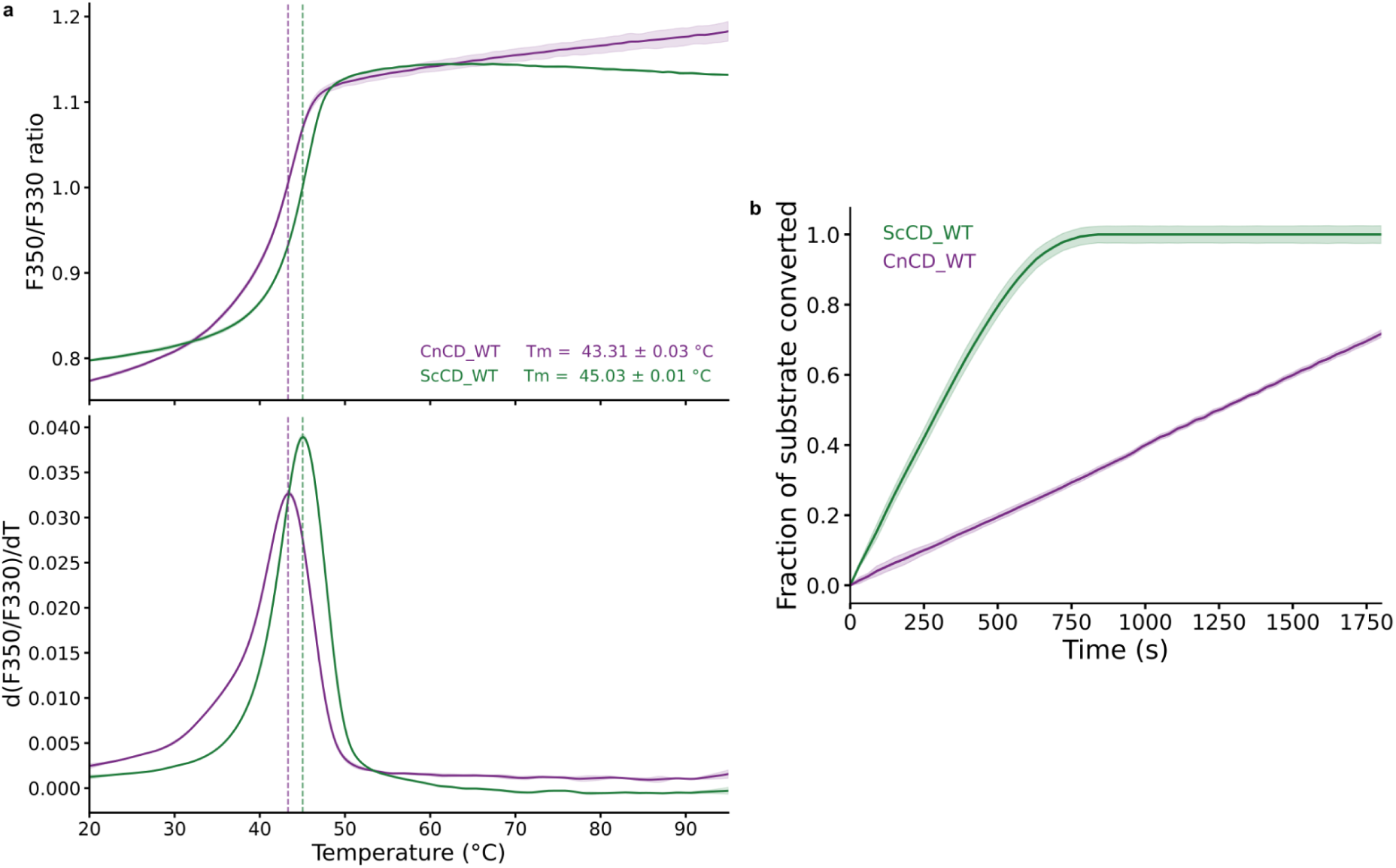
Thermal unfolding and time-dependent activity of CnCD and ScCD wild-type proteins. Thermal stability of wild-type proteins was assessed by nanoDSF. (**a**) The top panel shows the fluorescence ratio at 350 nm and 330 nm (F350/F330) as a function of temperature, and the bottom panel shows the first derivative of the fluorescence ratio with respect to temperature, d(F350/F330)/dT, used to determine T_m_. Curves represent the mean of three technical replicates (n = 3), and shaded regions indicate the standard deviation. Dashed lines indicate the corresponding T_m_ values in both panels. (**b**) Fraction of substrate converted as a function of time (s) for CnCD and ScCD WT enzymes. Substrate conversion was derived from the decrease in absorbance at 237 nm, which corresponds to 5-FC consumption. Raw absorbance values were corrected using blank controls and baseline-adjusted to the initial time point (t = 0 s). Replicate traces (n=3) were averaged and normalized to the total signal change observed for WT over the course of the reaction. All variants were scaled relative to WT, such that WT reaches a value of 1 at the final time point. Curves represent mean fractional 5-FC consumption relative to WT, and shaded regions indicate ± standard deviation.

**Supplementary Figure 12.**
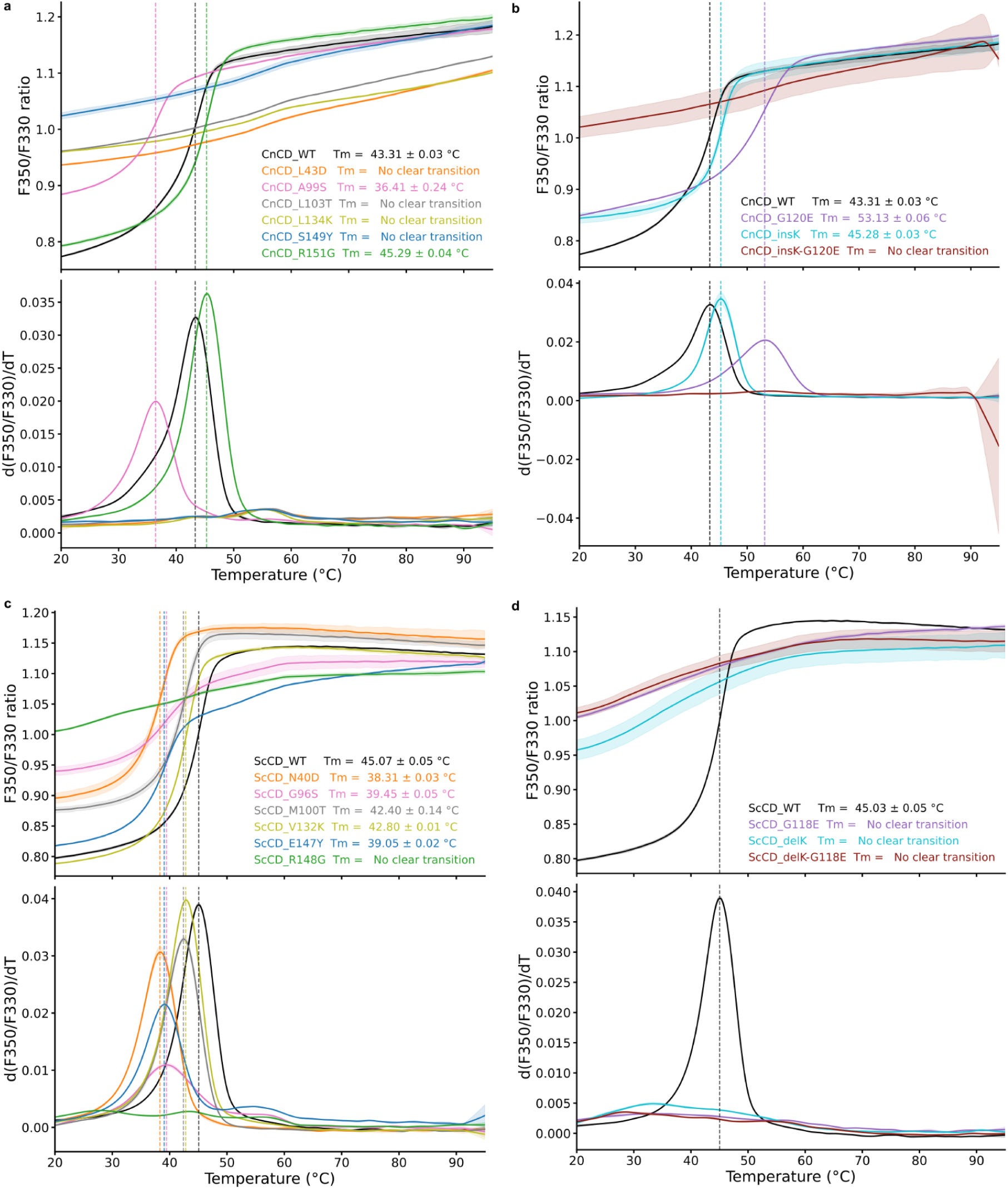
Thermal unfolding profiles of CnCD and ScCD variants. Thermal stability of variants was assessed by nanoDSF. The top panels show the fluorescence ratio at 350 nm and 330 nm (F350/F330) as a function of temperature, and the bottom panels show the first derivative of the fluorescence ratio with respect to temperature, d(F350/F330)/dT, used to determine T_m_. Curves represent the mean of three technical replicates (n = 3), and shaded regions indicate the standard deviation. Dashed lines indicate the corresponding T_m_ values in both panels. (**a**) CnCD variants. (**b**) CnCD variants carrying the G120E mutation and/or the insertion of a K residue between G119 and G120. (**c**) ScCD variants. (**d**) ScCD variants carrying the G118E mutation and/or the insertion of a K residue between K117 and G118.

**Supplementary Figure 13.**
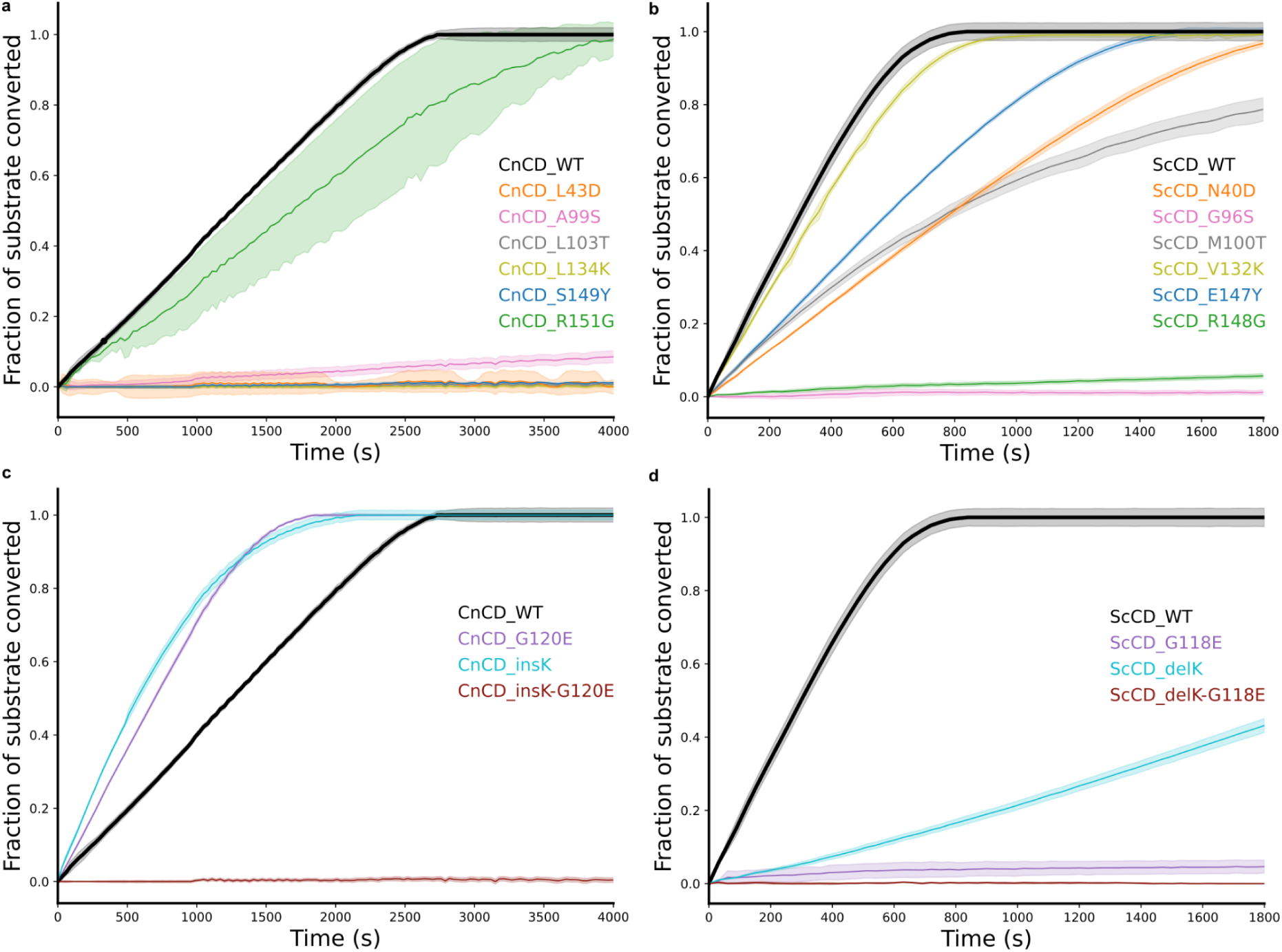
Time-dependent enzyme activity of CnCD and ScCD variants. Fraction of substrate converted as a function of time (s) for (**a**, **c**) CnCD and (**b**, **d**) ScCD WT enzymes and their variants. Substrate conversion was derived from the decrease in absorbance at 237 nm, which corresponds to 5-FC consumption. Raw absorbance values were corrected using blank controls and baseline-adjusted to the initial time point (t = 0 s). Replicate traces (n=3) were averaged and normalized to the total signal change observed for WT over the course of the reaction. All variants were scaled relative to WT, such that WT reaches a value of 1 at the final time point. Curves represent mean fractional 5-FC consumption relative to WT, and shaded regions indicate ± standard deviation.

**Supplementary Figure 14.**
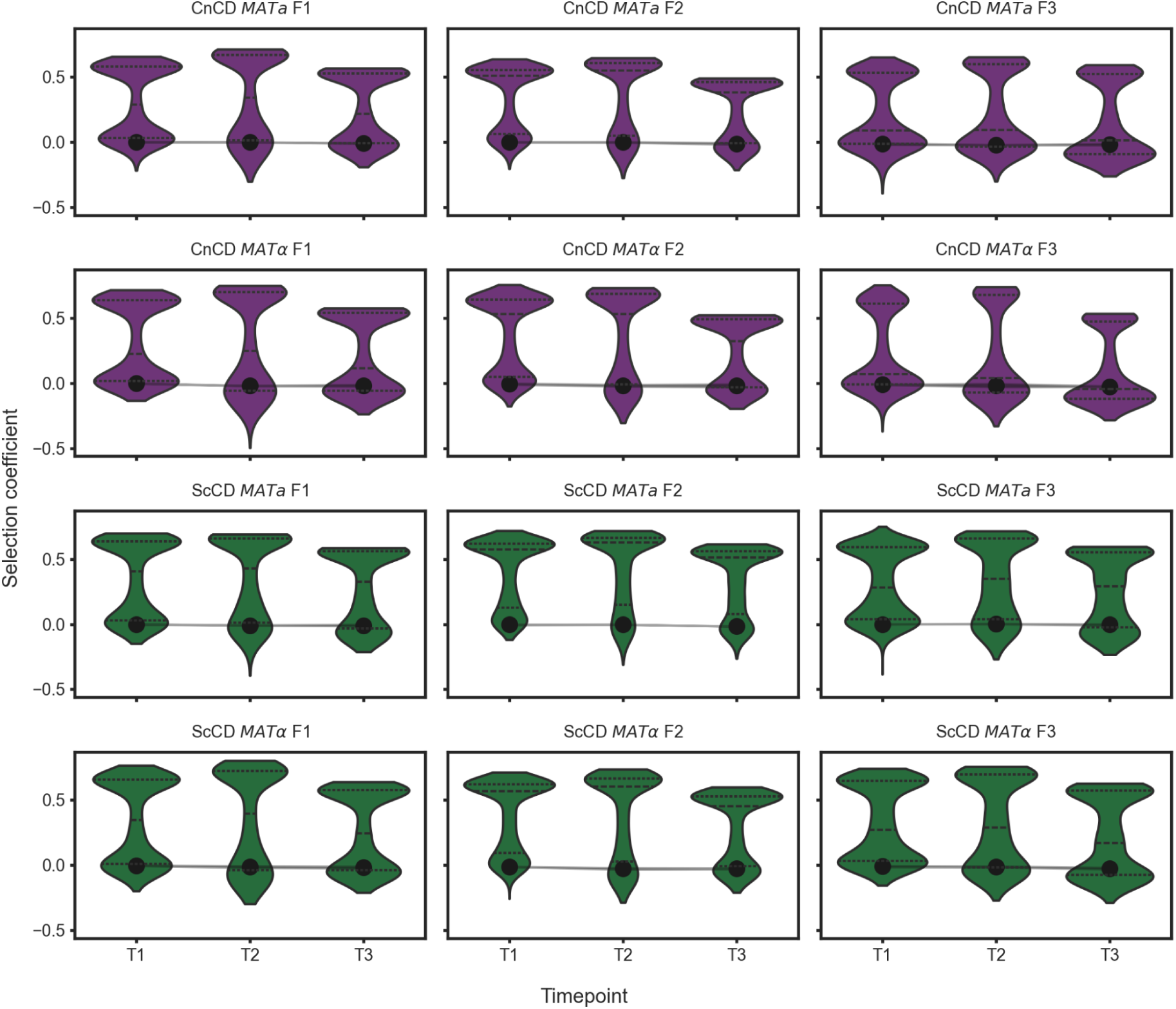
Selection coefficients at each time point. For each combination of species, mating type and fragment, the distribution of selection coefficients (average from two biological replicates) at each time point for single amino acid substitutions is shown as a violin plot. Values for the wild-type are indicated by black circles, with the band indicating the spread across biological replicates (2.5^th^ to 97.5^th^ percentile).

**Other Supplementary Materials for this manuscript include the following:**

Supplementary Tables 1 to 5 (provided as csv or excel spreadsheet).

